# Heat stress reveals the existence of a specialized variant of the pachytene checkpoint in meiosis of *Arabidopsis thaliana*

**DOI:** 10.1101/2021.06.28.450228

**Authors:** Joke De Jaeger-Braet, Linda Krause, Anika Buchholz, Arp Schnittger

## Abstract

Plant growth and fertility strongly depend on environmental conditions such as temperature. Remarkably, temperature also influences meiotic recombination and thus, the current climate change will affect the genetic make-up of plants. To further understand temperature effects on meiosis, we have followed male meiocytes of *Arabidopsis thaliana* by live cell imaging under three different temperature regimes, at 21°C and at heat shock conditions of 30°C and 34°C as well as after an acclimatization phase of one week at 30°C. This work led to a cytological framework of meiotic progression at elevated temperature. We found that an increase to 30°C, sped up meiotic progression with specific phases being more amenable to heat than others. An acclimatization phase often moderated this effect. A sudden increase to 34°C promoted a faster progression of meiosis in early prophase compared to 21°C. However, the phase in which cross-overs maturate was found to be prolonged at 34°C. Interestingly, mutants involved in the recombination pathway did not show the extension of this phase at 34°C demonstrating that the delay is recombination dependent. Further analysis revealed the involvement of the ATM kinase in this prolongation indicating the existence of a specialized variant of the pachytene checkpoint in plants.

## INTRODUCTION

The ambient temperature is one of the key environmental parameters that determines plant growth and fertility and has been the focal interest of many plant researchers. Understanding the plant response to temperature is further boosted by the ongoing climate change (Anderson et al., 2016; Collins, 2014; Couteau et al., 1999), during which crops are expected to be exposed to very high temperatures in the near future, threatening a sharp reduction in crop yield (Hatfield and Prueger, 2015; Yue et al., 2019). For example, a yield decrease of up to 22% for corn (*Zea mays*) could be observed with a 1°C increase in temperature (Kukal and Irmak, 2018). To rebuttal these detrimental effects and adjust breeding programs, it is vital to understand the changes temperature stress imposes on yield-related traits at the cellular and molecular level.

Central for sexual reproduction and fertility is meiosis and previous work has demonstrated its sensitivity to changes in environmental conditions, especially temperature as reviewed by (Bomblies et al., 2015; Morgan et al., 2017). Meiosis is a specialized cell division in which DNA replication is followed by two rounds of chromosome segregation (meiosis I and meiosis II), resulting in the reduction of DNA content by half as a prerequisite for a subsequent fusion of gametes and restoration of the full genome size. Furthermore, meiosis I plays an important role for the generation of genetic diversity via cross-over (CO) formation during prophase I and the new assortment of chromosome sets. COs are not only important for the generation of new allelic combinations but also ensure physical connections between homologous chromosomes (homologs) that are needed for their balanced segregation. COs are visible as chiasmata, that connect two homologs in the form of a bivalent.

The control and execution of meiotic recombination is highly conserved and like in other eukaryotes, meiotic recombination in plants is initiated by a conserved topoisomerase-like protein SPORULATION 11-1 (SPO11-1), and together with associated proteins catalyzes DNA double stranded breaks (DSBs) in early meiosis (Grelon et al., 2001; Hartung et al., 2007; Keeney et al., 1997; Stacey et al., 2006). Subsequently, DSBs are processed by the MRE11, RAD50 and NBS1 (MRN) protein complex and recognized by the recombinases DISRUPTED MEIOTIC cDNA1 (DMC1) (Bishop et al., 1992; Couteau et al., 1999) and RECA HOMOLOG RADIATION SENSITIVE 51 (RAD51) (Jachymczyk et al., 1981; Li et al., 2004). They mediate the invasion of the processed single stranded DNA into the DNA double strand of the homolog. In the absence of DMC1, DSBs are repaired by inter-sister recombination resulting in the absence of COs and hence, causing the formation of unconnected homologs, called univalents. In *rad51* mutants, DSBs are not repaired resulting in severely fragmented chromosomes and complete sterility of the mutant plants.

Towards the end of prophase I, all DSBs are resolved into either non-crossovers (NCOs) or COs. COs fall in one of the two classes: Type I COs rely on the ZMM proteins (acronym from the *Saccharomyces cerevisiae* Zip, Mer and Msh proteins), including MUTS HOMOLOG 4 (MSH4), and are formed at a distance to each other (CO interference) (Higgins et al., 2004; Su and Modrich, 1986). Type II COs are catalyzed by a protein called MMS AND UV SENSITIVE 81 (MUS81) and are not subjected to interference (Berchowitz et al., 2007; Interthal and Heyer, 2000). Homologs remain connected to each other until cohesin, a proteinaceous ring that embraces the sister chromatids of each homolog, is opened by cleavage of its alpha kleisin subunit, RECOMBINANT PROTEIN (REC8), along the chromosome arms in anaphase I paving the road for the separation of the homologs, each to opposite cell poles and hence, completing meiosis I (Cai et al., 2003).

The successful execution of meiotic recombination as means to equally segregate homologs and ensure genetic diversity is controlled by the pachytene checkpoint or meiotic recombination checkpoint in animals and yeast (Roeder and Bailis, 2000). This checkpoint delays meiotic progression until recombination defects are resolved. Consequently, several mutants, especially in the recombination pathway, *e.g*. *dmc1* mutants, trigger this checkpoint and a prolonged arrest, which can even lead to apoptosis in several species, including mouse (Barchi et al., 2005; Bishop et al., 1992; Lange et al., 2011; Rockmill et al., 1995).

A master regulator of the pachytene checkpoint is ATAXIA TELANGIECTASIA MUTATED (ATM), a kinase activated by DNA damage, that triggers checkpoint signaling, promotes DSB repair and in addition, controls the DSB number by regulating SPO11-1 activity via a negative feedback loop (Lange et al., 2011). While ATM is present in plants and fulfills several important functions during meiosis, it was thought so far that a pachytene checkpoint in plants did not exist since mutants, like *dmc1,* do not arrest at pachytene and instead complete meiosis leading to aneuploid gametes (Caryl et al., 2003; Couteau et al., 1999; Jackson et al., 2006; Jones and Franklin, 2008; Muyt et al., 2009).

Meiosis and in particular meiotic recombination are highly sensitive to environmental conditions, leading to meiotic failure in many different organisms, such as *Caenorhabditis elegans* (Bilgir et al., 2013), mouse (Nebel and Hackett, 1961), wheat (Pao and Li, 1948) and rose (Pecrix et al., 2011). Elevated temperatures affect the meiotic microtubule cytoskeleton, resulting in irregular spindle orientation, aberrant cytokinesis and the production of unreduced gametes, polyads and micronuclei in *Populus*, Rosa and *Arabidopsis thaliana* (Arabidopsis) (De Storme and Geelen, 2020; Hedhly et al., 2020; Pecrix et al., 2011; Wang et al., 2017).

Further, although the DSB numbers are reported to be unaffected at elevated temperatures in several organisms, *e.g*. yeast and Arabidopsis (Brown et al., 2020; Modliszewski et al., 2018), other aspects of the recombination pathway were found to be altered by temperature leading to diverse effects, which differ depending on the environmental conditions and species. Chiasma frequency was shown to be highly sensitive to environmental conditions. Upon high temperatures, the chiasma frequency was reduced in some species, such as female barley, *Tradescantia bracteata*, *Uvularia perfoliata* and wild garlic (Dowrick, 1957; Lloyd et al., 2018; Loidl, 1989; Modliszewski et al., 2018; Phillips et al., 2015), while it increased in other species, for instance in male barley, Arabidopsis and *Sordaria fimicola* (Lamb, 1969; Lloyd et al., 2018; Modliszewski et al., 2018; Phillips et al., 2015). In Arabidopsis, it was shown that the increase in CO frequency upon high temperatures is regulated via the Type I CO pathway (Lloyd et al., 2018; Modliszewski et al., 2018). In addition, CO distribution is also altered by heat stress (Dowrick, 1957; Higgins et al., 2012). In barley, high temperatures cause an increase in chiasmata at the interstitial/proximal region of chromosomes but an overall decrease in chiasmata per cell (Higgins et al., 2012). At very high temperatures, in many species, such as wheat, barley, wild garlic and *Cynops pyrrhogaster*, synapsis of the homologs fails resulting in the formation of univalents (Higgins et al., 2012; Loidl, 1989; Pao and Li, 1948; Yazawa et al., 2003).

To obtain further insights into temperature effects on meiosis, we followed in this study Arabidopsis male meiocytes under three different temperature regimes via live cell imaging. This led to a detailed picture of the meiotic progression under heat stress. A key discovery was that the length of pachytene/diakinesis is prolonged at 34°C, while in general, meiocytes progressed through meiosis much faster than at 21°C. An extension of pachytene/diakinesis was not observed when recombination was abolished. Since this extension was also eradicated in *atm* mutants, we conclude that Arabidopsis and likely other plants have a specialized form of the pachytene checkpoint that is only triggered by recombination intermediates but not the complete absence of recombination.

## RESULTS

### A cytological sensor of heat stress in meiocytes

To analyze the effects of increased temperatures on meiosis, we decided to apply three different heat conditions reflecting possible environmental stress scenarios in the future and matching conditions used in previous studies. Arabidopsis is typically grown between 18 and 24°C (our standard growth conditions being 21°C). As a first stress condition, we used an immediately applied heat shock of 30°C (HS30°C). In parallel, we allowed plants to acclimatize to 30°C for one week (long-term, LT30°C) before analyzing meiosis. The third condition was a greater heat stress of 34°C (HS34°C) that was also applied immediately.

However, the proper and reliable application of heat stress to multicellular structures, such as anthers, can be challenging when the focus is on particular cells, like meiocytes, which are surrounded by many different cell layers, such as the tapetum layer and the epidermis. The multicellular environment and the size of these structures have the capacity to buffer temperatures, hence making it difficult to exactly time the moment when the heat stress will reach the cells of interest.

To approach this problem, we made use of the observation that stress granules (SGs) are formed at elevated temperatures in different plant tissues, *e.g*. roots, leaves and hypocotyls (Chodasiewicz et al., 2020; Dubiel et al., 2020; Hamada et al., 2018; Kosmacz et al., 2019; Modliszewski et al., 2018). It was previously shown that these SGs in Arabidopsis seedlings contain the cell cycle regulator CYCLIN-DEPENDENT KINASE A;1 (CDKA;1)(Kosmacz et al., 2019). CDKA;1 is a major regulator of meiotic progression as well as recombination and is highly expressed in male meiocytes of Arabidopsis (Bulankova et al., 2010; Dissmeyer et al., 2007; Sofroni et al., 2020; Wijnker et al., 2019; Yang et al., 2020; Zhao et al., 2017; Zhao et al., 2012). To test whether CDKA;1 would change its homogenous cytosolic and nuclear localization pattern in meiosis upon heat stress, we applied different temperature regimes to male meiocytes from plants carrying the *CDKA;1:mVenus* and the *TagRFP:TUA5* reporters, and followed the localization pattern of CDKA;1:mVenus during meiosis (Sofroni et al., 2020).

Under our standard Arabidopsis growth conditions (21°C) and consistent with previous analyses, CDKA;1:mVenus is uniformly localized in both the cytoplasm and the nucleus. The localization shifts from preferential cytosolic to predominantly nuclear in late leptotene till early pachytene followed by increased cytosolic accumulation in pachytene and diakinesis. After anaphase I and anaphase II, CDKA;1:mVenus accumulates again in the reforming nuclei (Yang et al., 2020)(**Figure 1A**).

**Figure 1.**
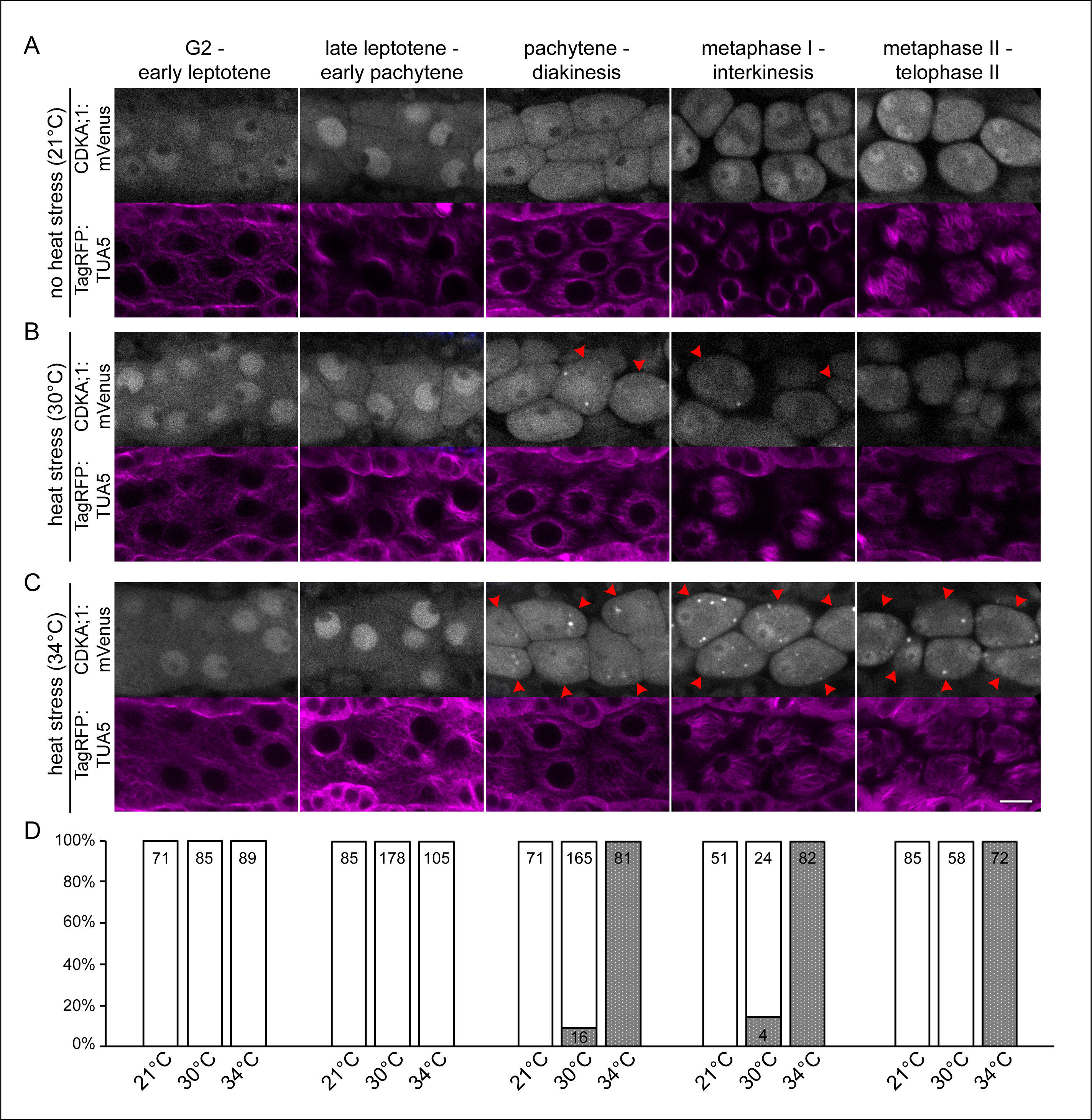
Localization of CDKA;1 in male meiocytes in control and stress conditions. CDKA;1:mVenus (first row; white) and TagRFP:TUA5 (second row; magenta) localization at control conditions of 21°C (A), heat stress of 30°C (B) and of 34°C (C) at different meiotic stages: G2-early leptotene (column 1), late leptotene-early pachytene (column 2), pachytene-diakinesis (column 3), metaphase I-interkinesis (column 4) and metaphase II-telophase II (column 5). Red arrowheads highlight cells with CDKA;1 localization at SGs. Scale bar: 10 um. Quantification of CDKA;1 SG formation on cellular level (D) per stage in percent; white bar: cells without SGs; grey bar: cells having at least one SG; the absolute sample size is given in the corresponding bar.

At elevated temperatures, 30°C and 34°C, we found the same cytosolic-nuclear localization dynamics of CDKA;1 (**Figure 1B,C**). At 34°C, no SGs were observed in early meiotic stages (n=0/89 in G2-early leptotene; n=0/105 from late leptotene till early pachytene), when CDKA;1 is preferentially localized to the nucleus of meiocytes (**Figure 1C,D**). Notably, SGs were readily formed at 34°C in all meiocytes from pachytene till diakinesis (n=81/81), from metaphase I till interkinesis (n=82/82), and from metaphase II till telophase II (n=72/72), *i.e.* the period when CDKA;1 starts to locate predominantly in the cytoplasm. These granules were immediately visible after the heat stress was applied, *i.e.* after 15 min, the time required to set up the acquisition for live cell imaging at the microscope. Thus, the formation of SGs occurs within the first 15 min of heat stress.

In contrast, CDKA;1 granules were rarely formed at 30°C, *i.e.* in only 9% and 14% of the meiocytes in pachytene/diakinesis (n=16/165) and from metaphase I till interkinesis (n=4/24), respectively (**Figure 1B,D**). In addition, the number of SGs per meiocyte was also lower at 30°C compared to granule-containing meiocytes at 34°C. Taken together, monitoring of SG formation allows a visual discrimination between 30°C and 34°C consistent with the previous observation that the temperature threshold for the formation of SGs is 34°C (Hamada et al., 2018). Importantly, this optical marker indicated that the ambient temperature reaches meiocytes in short time, *i.e.* less than 15 min, paving the road for the faithful application of different heat treatments and comparison by live cell imaging.

### Heat stress affects microtubule configurations in meiosis in a quantitative but not qualitative manner

After having confirmed that the applied temperature regime reached male meiocytes fast and faithfully, we turned towards addressing the general aim of this study, *i.e*. the question how increased temperature affects the dynamics of meiosis. To address this, we aimed to use a previously established live cell imaging method for meiosis (Prusicki et al., 2019). A crucial finding in this approach was the observation that meiosis can be dissected by so-called landmarks that occur in a predictable order and that reflect highly defined cytological stages, for instance using fluorescently labeled microtubules (MT, TagRFP:TUA5). Thus, these landmarks not only allow the staging of meiocytes but also provide a mean to reveal the dynamics of meiosis by determining the time between landmarks.

In brief, MT have the following dynamics during male meiosis: During G2-early leptotene, MT are first homogenously distributed in meiocytes with the nucleus in the center, called MT array state 1 (**Supplemental Figure 1A**). Then, they will gradually polarize into a half moon-like structure on one side of the nucleus, MT array state 2-3-4, from late leptotene till early pachytene (**Figure 2A, Supplemental Figure 1B**). This develops further into a full moon-like structure entirely surrounding the nucleus, MT array state 5-6, during pachytene, diplotene and diakinesis (**Figure 2B, Supplemental Figure 1C**). After nuclear envelope breakdown (NEB), the pre-spindle transforms into the first meiotic spindle, MT array state 7-8-9, from metaphase I till anaphase I (**Figure 2C, Supplemental Figure 1D**). Next, MT reorganize around the two newly formed nuclei and central MT form a phragmoplast-like structure, MT array state 10-11, at telophase I and interkinesis (**Figure 2D, Supplemental Figure 1E**). The second division is characterized by the formation of two pre-spindles, followed by two spindles, MT array state 12-13, from metaphase II till anaphase II (**Figure 2E, Supplemental Figure 1F**). Phragmoplast-like structures, MT array state 14 are visible at telophase II (**Figure 2F, Supplemental Figure 1G**) until cytokinesis, resulting in tetrads, the four meiotic products.

**Figure 2.**
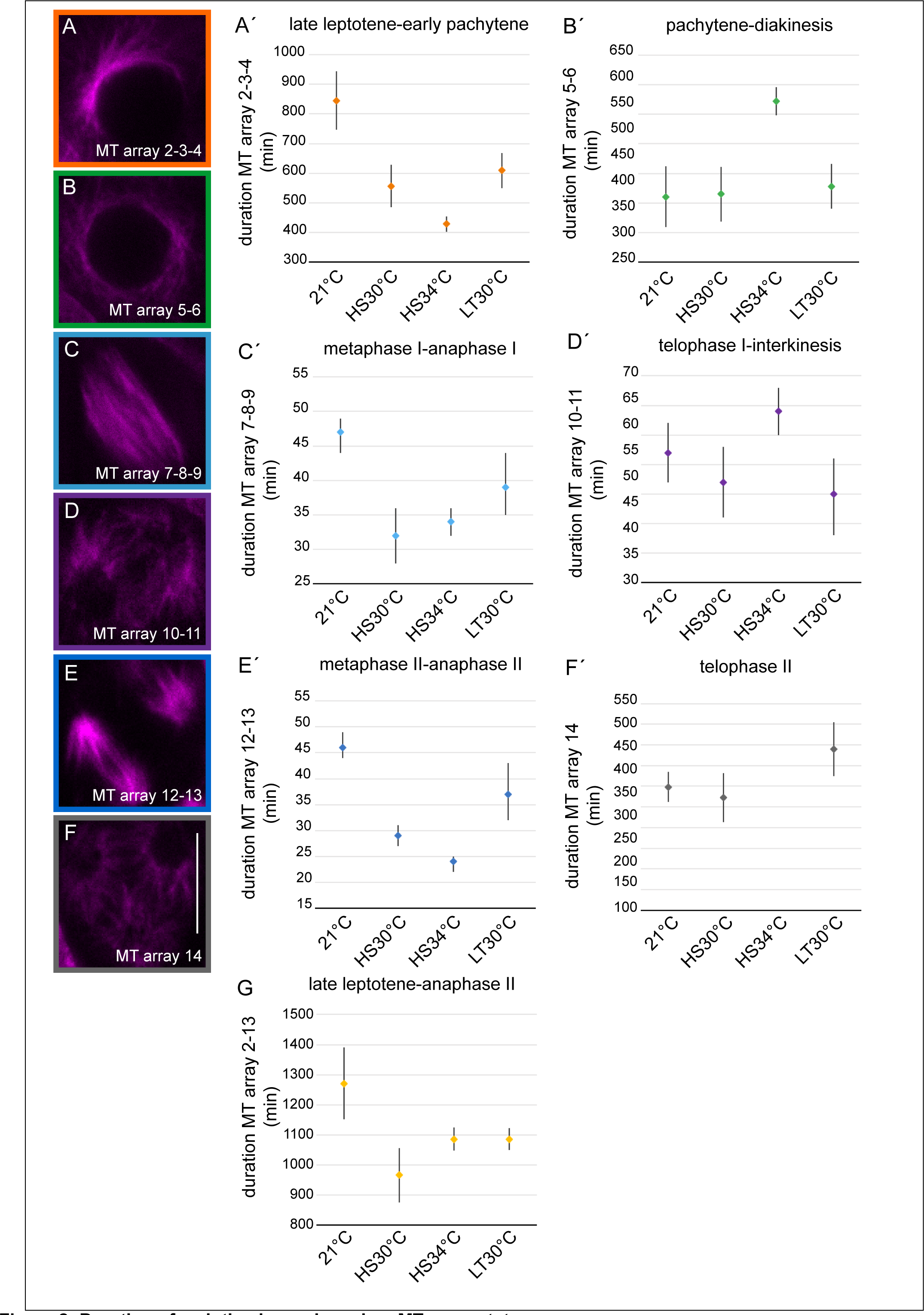
Duration of meiotic phases based on MT array states. Confocal images of the MT array states (A-F) and the corresponding predicted median times (in min) with 95% confidence inter- vals in control (21°C) and heat conditions (HS30°C, HS34°C and LT30°C) (Á-F′); (A,A ′; orange) MT array state 2-3-4, late lepto- tene-early pachytene; (B,B′; green) MT array state 5-6, pachytene-diakinesis; (C,Ć; light blue) MT array state 7-8-9, metaphase I-anaphase I; (D,D′; purple) MT array state 10-11, telophase I-interkinesis; (E,É; dark blue) MT array state 12-13, metaphase II-anaphase II; (F,F′; grey) MT array state 14, telophase II. Scale bar: 10 um. Predicted median time (in min) of MT array states 2-13 with the 95% confidence interval in control (21°C) and heat conditions (HS30°C, HS34°C and LT30°C) (G; yellow).

Analyzing meiosis at HS30°C, HS34°C and LT30°C, we confirmed that meiosis does not arrest upon exposure to these temperature regimes, consistent with previous studies (De Storme and Geelen, 2020; Lei et al., 2020). Importantly, in all movies taken at higher temperature (in total 46), the meiocytes progressed through the same MT array states as previously found for 21°C (**Supplemental Figure 1, Supplemental Movies 1-4**, (Prusicki et al., 2019)).

MT stability and polymerization are known to be temperature-sensitive, (Bannigan et al., 2007; Li et al., 2009a; Liu et al., 2017; Song et al., 2020; Wu et al., 2010). Consistently, we observed quantitative changes in some MT structures confirming that meiocytes were exposed to elevated temperatures. As revealed by pixel intensity quantification of meiocytes in MT array state 6, in which MT are fully surrounding the nucleus (**Figure 3A,B**), we found that the measured intensity of the fluorescence signal of TagRFP:TUA5 dropped upon both HS30°C and HS34°C in comparison to 21°C, indicating that MT density is reduced. Notably, this reduction is reverted at LT30°C, implying the existence of an adaptation mechanism for meiosis to heat. In addition, irregular spindle structures were observed at 34°C but not at lower temperatures (**Figure 3C**), consistent with previous analyses (De Storme and Geelen, 2020; Lei et al., 2020).

**Figure 3.**
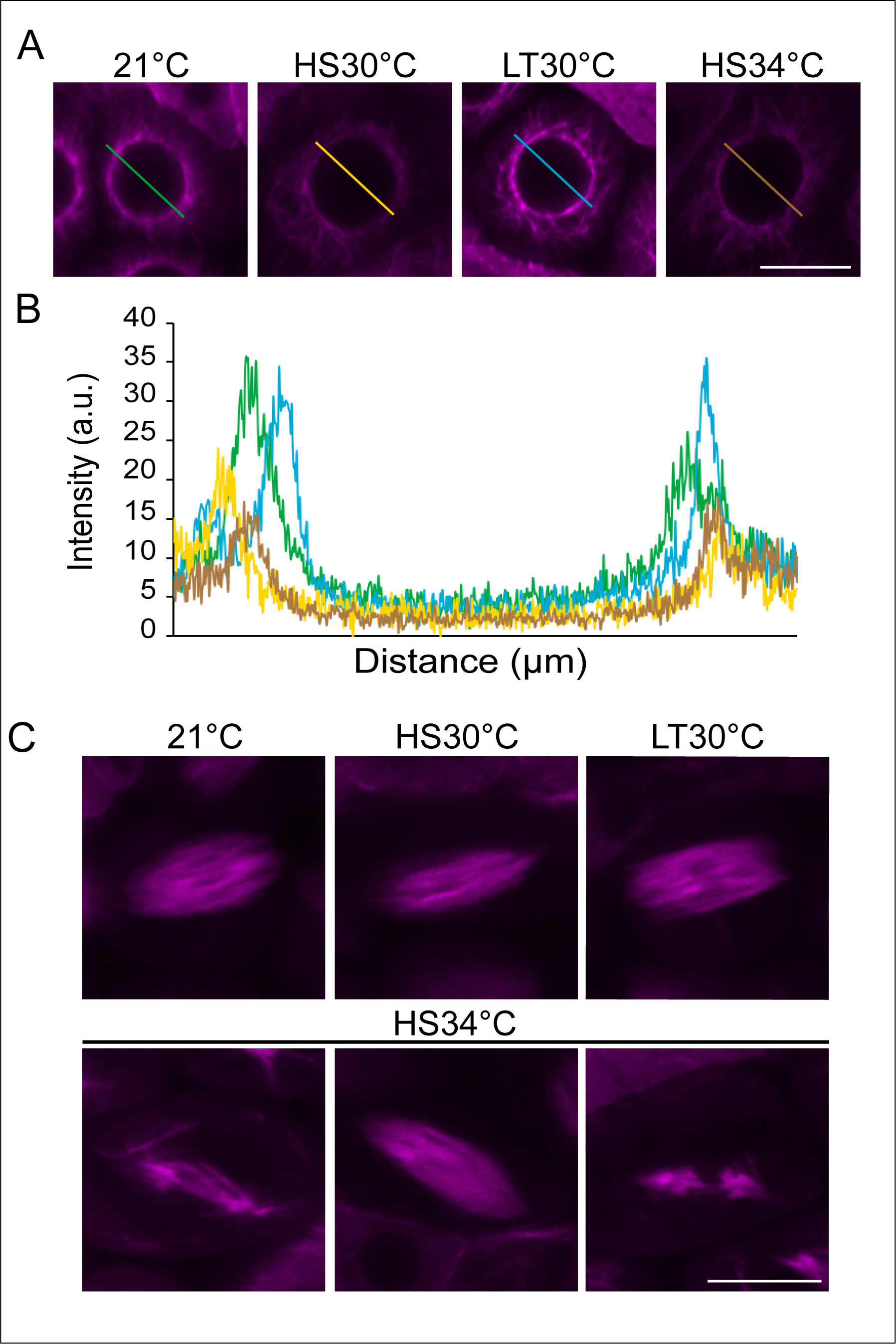
Microtubule array in wildtype in control and heat stress conditions. Confocal images of meiocytes expressing TagRFP:TUA5 (magenta) at MT array state 6 in control conditions (21°C), heat shock conditions (HS30°C, LT30°C and HS34°C) (A). Pixel intensity plot of a section crossing through the middle of the cell (distance in um) in MT array state 6 in 21°C (green), HS30 °C (yellow), LT30°C (blue) and HS34°C (brown) (B), section lines also highlighted in (A). Confocal images of meiocytes expressing TagRFP:TUA5 (magenta) at MT state 8-9 in 21°C, HS30°C, LT30°C and HS34°C (C). Scale bar: 10 um.

Taken together, the quantitative but not qualitative changes of the typical meiotic MT configurations allow using the adoption of characteristic MT arrays for staging of meiosis during live cell imaging. At the same time, the quantitative effects on the MT arrays corroborate the previous finding that meiocytes successfully received the heat treatment in our experimental set up.

### Duration of meiosis under heat stress

The next challenge to overcome for the evaluation of meiotic progression at elevated temperatures was the question how the MT-based dissection of the different heat stress experiments could be statistically compared with the control growth conditions. This is not a trivial question since the analyses of meiocytes within one anther-sac cannot be regarded as statistically independent measurements but represent clustered data. In addition, the above-mentioned nature of defined meiotic stages gives rise to a multi-state nature of our dataset. Moreover, our measurements occasionally did not capture the exact start and/or end point (left, right and/or interval censored data) of a MT array state since the observed anthers sometimes move out the focal plane (but also occasionally move into focus again).

The combination of the three characteristics of our data, *i.e*. clustering data, left/right and/or interval censoring, as well as having a multistate nature, in one statistical model was not possible. Therefore, we reduced the complexity of the analysis and built one separate model for each state. This also allowed us to simplify the mixture of left/right and/or interval censored data with respect to the duration of each state to interval (and right) censoring.

We applied parametric models for interval-censored survival time data with a clustered sandwich estimator of variance to address the clustering of meiocytes within anther-sacs, including effects of the heat treatment, genotype and their interactions. The underlying distribution of the parametric model was chosen based on the Akaike Information Criterion (AIC) with exponential, Gompertz, log-logistic, Weibull and log-normal distribution as candidates.

The models used information from all cells which had at least one observation in the respective stage. The event of interest is the transition of a cell from one stage to the next. Each cell for which the exact beginning and end of the stage were known was modelled as having an event with the event time as the difference between the start of the next stage and the end of the previous stage. Cells where the exact time points of either the transition from the previous stage to the stage of interest or to the next stage were not known were modelled as interval-censored data points with the lower limit of the interval being the time where the cell was observed in this specific state and the upper limit of the interval being one time unit before when the cell was observed in the previous or next stage, respectively. If for a cell a certain stage either at the beginning or end was not observed the cell was modelled as right-censored with the censoring time being the minimum observed time for this cell in the stage of interest.

With the imaging and evaluation system in hand, we then addressed what the effect of HS30°C, HS34°C and LT30°C has on the total length of meiosis. To this end, plants were grown under long-day conditions in highly controlled growth chambers (see material and methods). For the long-term heat treatment, plants were transferred to constant 30°C around bolting time. At flowering stage, flower buds were dissected for live cell imaging of male meiocytes, as described in (Prusicki et al., 2020). For the heat shock treatments, only flower buds which were in MT array state 1 were used for the prediction of the duration of the different meiotic states upon heat shock. The determination of the meiotic duration relies on defined start and end points of MT states (called events). Since this is not possible for MT state 1 (no start point), the first stage that could be temporally evaluated was MT state 2-3-4.

From a total of 59 movies, we first aimed for movies that covered all MT array states (2-14) under the four temperature regimes. Unfortunately for HS34°C, we were not able to reliably determine the end point of MT array state 14 as the fluorescent signal of the MT became very poor, possibly due the fact that MT are more defusedly organized at high temperature versus control conditions (described above, **Figure 3**) and photo-bleaching after long time lapses.

To compare the overall meiotic duration at all heat conditions, we excluded MT array state 14 for this analysis and only considered movies capturing MT array states 2-13 (23 movies). Then, we built a separate parametric model for the complete duration (as described above), resulting in the total predicted median time, together with the 95% confidence interval (CI).

The duration of MT array states 2-13 at 21°C was determined to have a predicted median time of 1271 min (*i.e*. 21.2 h, CI 1151-1390 min, **Figure 2G, Supplemental Table 1**), This value matched very well the previous analyses of the duration of male meiosis in Arabidopsis by pulse-chase experiments and live cell imaging, underlining the robustness of our analysis and the reproducibility of meiotic progression at 21°C (Armstrong et al., 2003; Prusicki et al., 2019; Sanchez-Moran et al., 2007; Stronghill et al., 2014).

Next, the duration of meiosis under the heat conditions was analyzed, resulting in a predicted median time of 966 min (*i.e*. 16.1 h, CI 876-1056 min) upon HS30°C, 1086 min (*i.e*. 18.1 h, CI1048-1124 min) upon HS34°C and 1086 min (*i.e*. 18.1 h, CI 1050-1122 min) upon LT30°C (**Figure 2G, Supplemental Table 1**). These data confirm previous observations that meiosis progresses faster under elevated temperatures in comparison to control conditions, also demonstrating that our experimental system can be faithfully used to study the effect of heat on meiosis (Bennett et al., 1972; Draeger and Moore, 2017; Stefani and Colonna, 1996; Wilson, 1959).

### Duration of individual meiotic phases under heat stress

The live cell imaging approach together with the model based estimation of the duration of the MT array states, allowed us then to target the main aim of this study, *i.e.* to obtain a detailed and meiotic-phase specific assessment of the meiotic progression under elevated temperatures.

At 21°C, a total of 206 meiocytes were observed and the predicted median time per MT array state was calculated from those cells for which we had at least one observation in that specific state (**Table 1, Supplemental Table 1**). The median time in MT array state 2-3-4 was predicted to be 845 min (CI 746-944 min, **Figure 2A′**). Followed by MT array state 5-6, which had a predicted median time of 360 min (CI 309-412 min, **Figure 2B′**), MT array state 7-8-9 takes 47 min (CI 44-49 min, **Figure 2C′**) and MT array state 10-11 spans 52 min (CI 47-57 min, **Figure 2D′**). After that, the second meiotic division follows with a predicted median time of 46 min for MT array state 12-13 (CI 44-49 min, **Figure 2E′**), finishing the meiotic division with 219 min for MT array state 14 (CI 205-234 min, **Figure 2F′**).

**Table 1.** Overview of the duration of the meiotic phases based on the MT array states. Predicted median times and 95% confidence intervals (in min) of MT array state 2-3-4 (late leptotene-early pachytene), MT array state 5-6 (pachytene- diakinesis), MT array state 7-8-9 (metaphase I- anaphase I), MT array state 10-11 (telophase I- interkinesis), MT array state 12-13 (metaphase II- anaphase II) and MT array state 14 (telophase II) of wildtype at 21°C, HS30°C, HS34°C, LT30°C; recombination mutants *spo11*, *dmc1* and *msh4* at 21°C and HS34°C and *atm* mutant at 21°C and HS34°C. (n= number of cells observed). NA: not analysed.

| MT array state | 2-3-4<br>late leptotene-early pachytene |  |  | 5-6<br>pachytene-diakinesis |  |  | 7-8-9<br>metaphase I-anaphase I |  |  | 10-11<br>telophase I-interkinesis |  |  | 12-13<br>metaphase II-anaphase II |  |  | 14<br>telophase II |  |  |
| --- | --- | --- | --- | --- | --- | --- | --- | --- | --- | --- | --- | --- | --- | --- | --- | --- | --- | --- |
| Meiotic stage |  |  |  |  |  |  |  |  |  |  |  |  |  |  |  |  |  |  |
| Predicted time<br>(in min) | Median | 95% Conf.<br>Interval |  | Median | 95% Conf.<br>Interval |  | Median | 95% Conf.<br>Interval |  | Median | 95% Conf.<br>Interval |  | Median | 95% Conf.<br>Interval |  | Median | 95% Conf.<br>Interval |  |
| Treatment (n) |  |  |  |  |  |  |  |  |  |  |  |  |  |  |  |  |  |  |
| 21°C<br>(206) | 845 | 746 | 944 | 360 | 309 | 412 | 47 | 44 | 49 | 52 | 47 | 57 | 46 | 44 | 49 | 219 | 205 | 234 |
| HS30°C<br>(133) | 556 | 485 | 628 | 365 | 319 | 411 | 32 | 28 | 36 | 47 | 41 | 53 | 29 | 27 | 31 | 209 | 185 | 233 |
| HS34°C<br>(188) | 428 | 403 | 453 | 522 | 498 | 546 | 34 | 32 | 36 | 59 | 55 | 63 | 24 | 22 | 25 | NA | NA | NA |
| LT30°C<br>(211) | 609 | 550 | 667 | 378 | 340 | 416 | 39 | 35 | 44 | 45 | 38 | 51 | 37 | 32 | 43 | 256 | 230 | 282 |
| *spo11* 21°C<br>(224) | 1119 | 1031 | 1206 | 374 | 349 | 399 | 72 | 67 | 76 | 63 | 58 | 67 | 48 | 45 | 52 | 356 | 326 | 385 |
| *dmc1* 21°C<br>(157) | 1056 | 929 | 1184 | 343 | 331 | 355 | 67 | 63 | 71 | 63 | 59 | 67 | 47 | 45 | 49 | 281 | 262 | 301 |
| *msh4* 21°C<br>(193) | 951 | 861 | 1040 | 314 | 299 | 329 | 67 | 63 | 71 | 59 | 56 | 63 | 49 | 46 | 52 | 274 | 253 | 294 |
| *spo11* HS34°C<br>(198) | 626 | 572 | 681 | 412 | 393 | 431 | 35 | 32 | 38 | 54 | 49 | 58 | 23 | 21 | 26 | NA | NA | NA |
| *dmc1* HS34°C<br>(160) | 565 | 526 | 605 | 383 | 362 | 403 | 30 | 28 | 32 | 57 | 54 | 60 | 22 | 21 | 23 | NA | NA | NA |
| *msh4* HS34°C<br>(116) | 571 | 536 | 606 | 398 | 346 | 450 | 32 | 30 | 34 | 49 | 43 | 55 | 24 | 22 | 26 | NA | NA | NA |
| *atm* 21°C<br>(228) | 834 | 761 | 908 | 295 | 270 | 321 | 45 | 42 | 49 | 60 | 55 | 66 | 43 | 40 | 46 | 245 | 230 | 260 |
| *atm* HS34°C<br>(172) | 702 | 640 | 764 | 350 | 330 | 370 | 31 | 29 | 33 | 55 | 50 | 60 | 26 | 23 | 28 | NA | NA | NA |

Next, male meiosis subjected to the three different temperature regimes was analyzed in the same way. A total of 133, 188 and 211 meiocytes were observed for HS30°C, HS34°C and LT30°C, respectively and the predicted median time per state was calculated from those cells for which we had at least one observation in that specific state (**Table 1, Supplemental Table 1**). The duration of MT array state 2-3-4 upon higher temperature was decreased compared to 21°C (845 min), with a predicted median time of 556 min upon HS30°C (CI 485-628 min), 428 min upon HS34°C (CI 403-453 min) and 609 min upon LT30°C (CI 550-667 min, **Figure 2A′**). This shows that the increase in temperature generally decreases the duration of this phase.

A strikingly different behavior was revealed for the next phase, *i.e.* MT array state 5-6. While upon exposure to HS30°C and LT30°C the predicted median time was 365 min (CI 319-411 min) and 378 min (CI 340-416 min), respectively, HS34°C resulted in a median of 522 min (CI 498-546 min, **Figure 2B′**). This was a much longer duration of this phase compared to 21°C (360 min), presenting a prolongation of ∼2.7 h.

After NEB, the meiocytes undergo the first round of chromosome segregation, *i.e*. MT array state 7-8-9, with a predicted median time of 32 min (CI 28-36 min) upon HS30°C, 34 min (CI 32-36 min) upon HS34°C and 39 min (CI 35-44 min) upon LT30°C, which is decreased compared to 21°C (47 min, **Figure 2C′**).

The following MT array state 10-11 spanned 47 min (CI 41-53 min) upon HS30°C, 59 min (CI 55-63 min) upon HS34°C and 45 min (CI 38-51 min) upon LT30°C (**Figure 2D′**). Whether the differences between HS34°C and 21°C (52 min) is biologically relevant is to be resolved.

Upon HS30°C, HS34°C and LT30°C, the second round of chromosome segregation, MT array state 12-13, spanned 29 min (CI 27-31 min, n=77), 24 min (CI 22-25 min, n=175) and 37 min (CI 32-43 min, n=150), respectively (**Figure 2E′**). Although statistically different compared to 21°C (46 min), the biological relevance is not clear at this moment and needs to be investigated in future.

The end of the meiotic division upon heat treatment was predicted using MT array state 14, which spanned 209 min (CI 185-233 min, n=65) upon HS30°C and 256 min (CI 230-282 min, n=144) upon LT30°C (**Figure 2F′**). The pairwise comparison of 21°C and LT30°C showed a statistical difference which needs to be investigated in future. Last, as described before, for HS34°C we were not able to predict the duration of MT array state 14.

### Exposure to high temperature causes chromosomal defects

To investigate the unexpected prolongation of late prophase at 34°C (**Figure 2B′**) in more detail, we first performed chromosome spreads from fixed flower buds at the different temperature regimes to investigate chromosomal behavior in our heat conditions and which could be confirmed in previous studies (De Storme and Geelen, 2020; Hedhly et al., 2020; Higgins et al., 2012).

At control growth conditions, decondensed chromatin becomes organized into chromosomes which will gradually condense during early prophase I and reach a fully paired state at pachytene. The paired homologs condense further, where chiasmata hold homologs together, finally reaching the highest condensed state at diakinesis with the formation of five bivalents that align at the metaphase plate during metaphase I (**Supplemental Figure 2A**). At both HS30°C and LT30°C, homologs condense and fully pair. Occasionally, two or more bivalents appear to be entangled at diakinesis and metaphase I forming chromosome bridges, suggesting interconnected non-homologous chromosomes. In addition, chromosome fragments and univalents were infrequently observed (**Supplemental Figure 2B,C**). In contrast to 21°C and 30°C, fully paired homologs could not be found at 34°C. Further, chromosome spreads of cells in diakinesis and metaphase I at 34°C revealed the formation of mainly 10 univalents. In addition, chromosome bridges were visible between both homologs and non-homologous chromosomes (**Supplemental Figure 2D**).

Thus, consistent with previous data, we find that high temperature causes recombination defects, which increase with rising temperatures (Bomblies et al., 2015; Brown et al., 2020; De Storme and Geelen, 2020; Higgins et al., 2012; Modliszewski et al., 2018; Morgan et al., 2017; Phillips et al., 2015).

### Synaptonemal complex formation is defective at 34°C

Given the central role of the formation of the chromosome axis for pairing and meiotic recombination, we next analyzed the localization of the previously generated reporters ASY1:RFP and ZYP1b:GFP upon 30°C and 34°C (Yang et al., 2019; Yang et al., 2020). ASYNAPTIC 1 (ASY1) is a chromosome axis-associated protein, which plays a major role in the initiation of synapsis and recombination (Armstrong et al., 2002; Caryl et al., 2000; Sanchez-Moran et al., 2007). ZIPPER 1 (ZYP1) encodes for a component of the transversal element of the synaptomenal complex (SC), which is thought to be crucial for the maturation of COs and CO interference (Capilla-Perez et al., 2021; France et al., 2021; Higgins et al., 2005; Osman et al., 2006).

At standard conditions, 21°C, ASY1 localizes to the chromosome axis from early leptotene till pachytene. During zygotene, when the formation of the SC is initiated, ASY1 gets largely depleted from the chromosome axis and ZYP1b signal starts to appear on chromosomes from where it gradually forms into a linear structure, resulting in the labeling of the complete chromosome axis at pachytene (**Figure 4A**).

**Figure 4.**
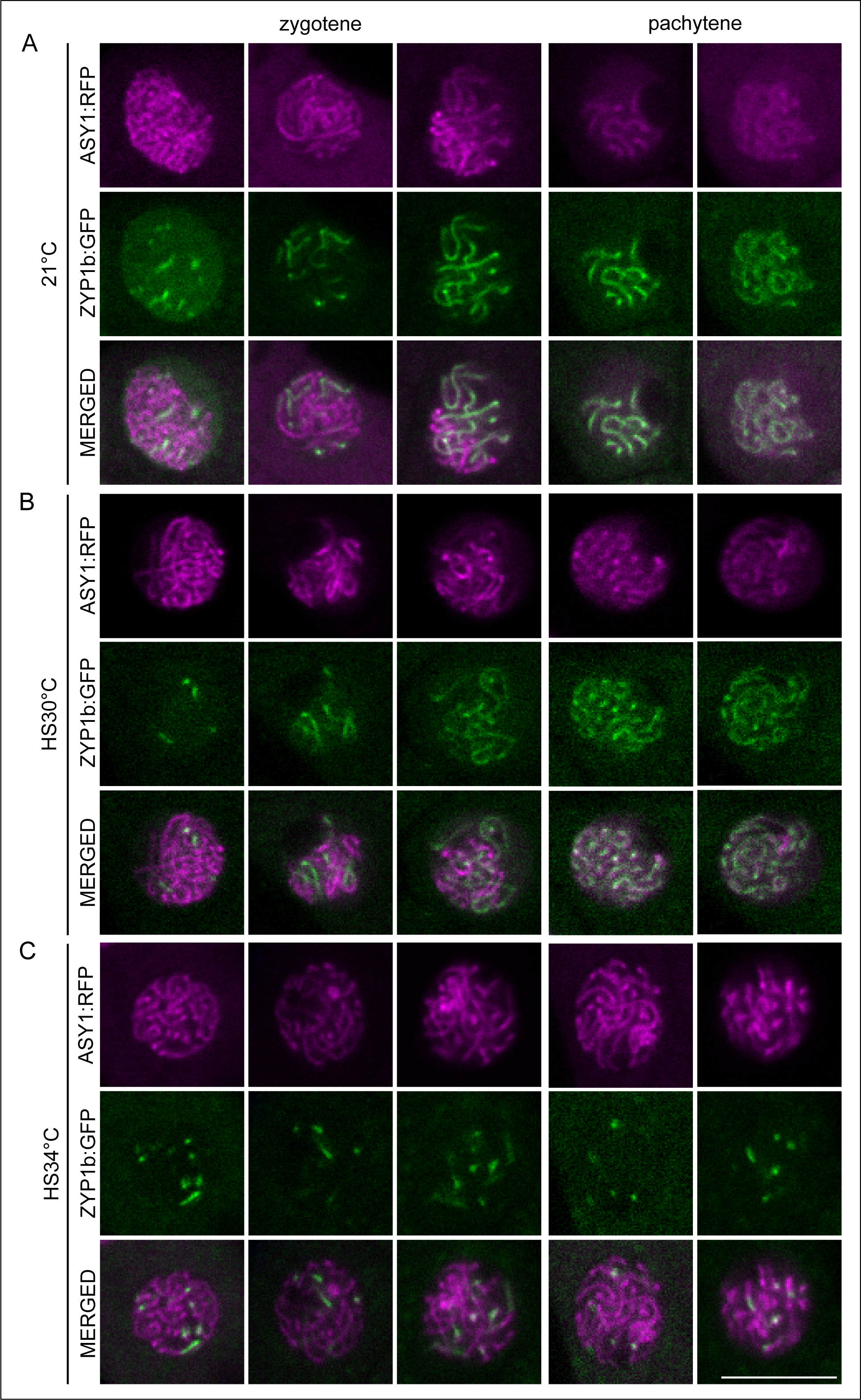
Synaptonemal complex elements ASY1 and ZYP1b localization upon heat stress. Confocal images of the nucleus of meiocytes at 21°C (A), HS30°C (B) and HS34°C (C) of SC elements ASY1:RFP (magenta, first row) and ZYP1b:GFP (green, second row) separately and merged (third row) at zygotene (columns 1-3) and pachytene (columns 4-5). Scale bar: 10 um.

Under high temperatures, 30°C and 34°C, the localization of ASY1 at the chromosome axis was unaffected and ZYP1b started to form short linear stretches at the chromosome axis during zygotene (**Figure 4B,C**). At 30°C, ZYP1b continues to label the full length of the axis, in contrast to 34°C, at which only small stretches of ZYP1b signal could be observed, suggesting that ZYP1b loading is initiated properly but discontinues (**Figure 4B,C**). This result is in accordance with previous findings showing that synapsis is obstructed upon very high temperature leading to the formation of abnormal structures, called polycomplexes (Bilgir et al., 2013; Higgins et al., 2012; Loidl, 1989).

### Late heat shock 34°C does not cause an elongation of pachytene/diakinesis

Seeing defective SC formation at 34°C, we asked whether events between zygotene and pachytene are particularly sensitive to heat stress and hence, responsible for the delay of NEB. Therefore, we specifically applied heat stress only from MT state 2-3-4 (zygotene) onward and compared the effect of this treatment with the previously applied heat shock before MT state 1, *i.e*. from pre-meiosis-leptotene onwards, by live cell imaging. Since we showed above that male meiocytes perceive heat stress in less than 15 min, we were confident that a late heat shock can allow distinguishing the temperature effects on early versus late prophase faithfully.

The predicted median time of MT array state 5-6 was calculated as described before and the comparison of flower buds in MT array state 1 and MT array state 2-3-4 at the onset of the heat stress was performed. We did not observe a statistical difference between early and late applied HS30°C, with a predicted median time of 313 min for MT array state 5-6 at late HS30°C (CI 270-355 min, **Figure 5A, Supplemental Table 1**). Remarkably, the extension of MT state 5-6 seen at early HS34°C (median time of 522 min) did not take place when we applied a late HS34°C, with a median time of 393 min (CI 349-437 min, **Figure 5B, Supplemental Table 1**). This suggests that the prolongation of MT state 5-6 is most probably not due to temperature effects on regulatory processes that take place at the moment of SC formation or on the SC itself. Rather early steps in prophase I, *e.g*. the initiation of meiotic recombination, might be temperature sensitive and subsequently affect the duration of pachytene/diakinesis.

**Figure 5.**
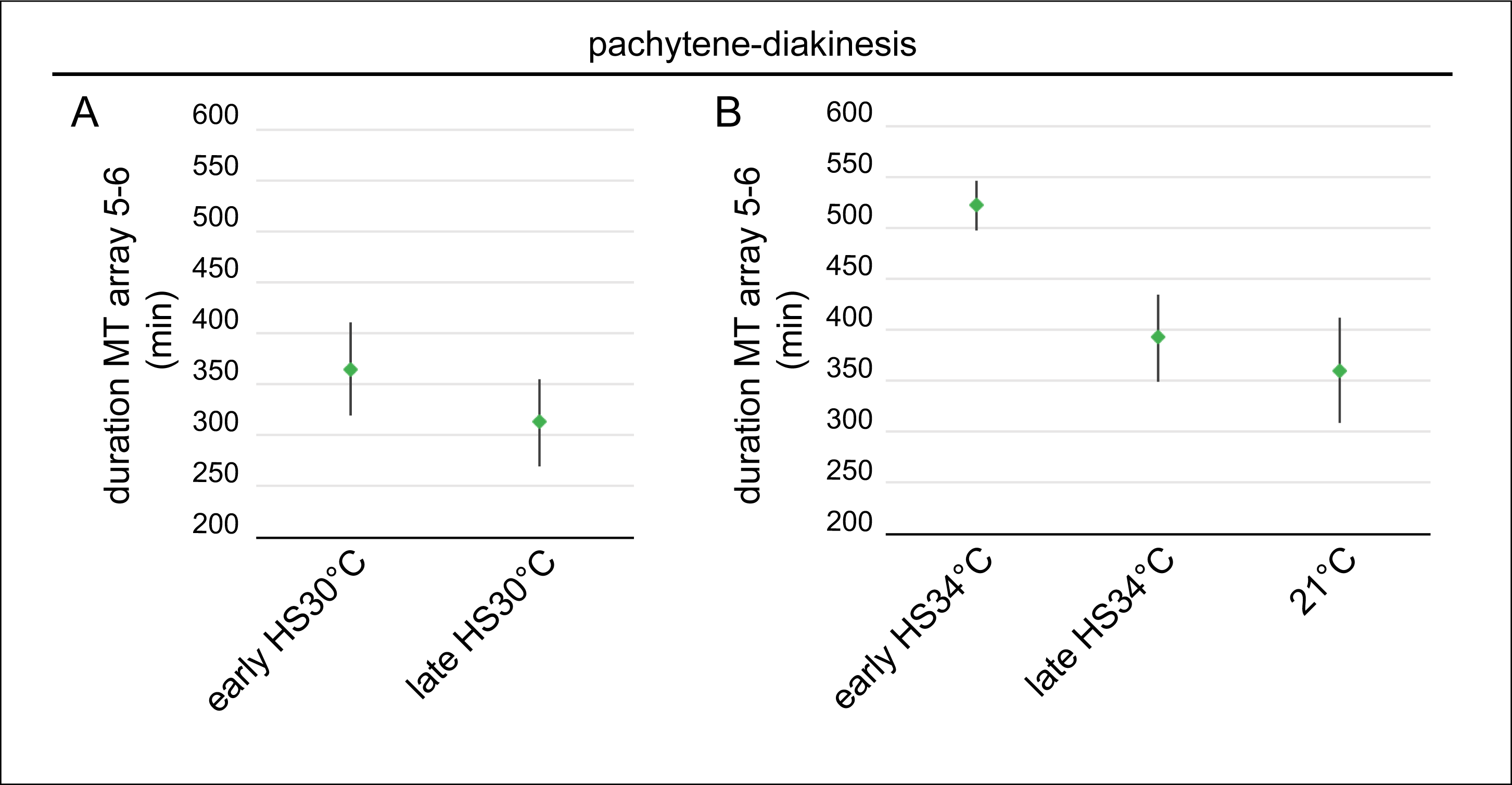
Effect of early and late heat shock on the duration of MT array state 5-6. The predicted median time (in min) with 95% confidence intervals of pachytene-diakinesis (MT array state 5-6; green) of early HS30°C versus late HS30°C (A) and early HS34°C versus late HS34°C and 21°C (B). Early HS30°C, early HS34°C and 21°C as shown in Figure 2B′.

### Loss of recombination *per se* does not cause the elongation of pachytene/diakinesis

To address to what degree a failure of recombination, which is initiated in early prophase, causes a pachytene/diakinesis delay, we first made use of the well-characterized *spo11-1* mutant (Grelon et al., 2001; Hartung et al., 2007), in which recombination is completely abolished due to a failure to form DSBs. The *TagRFP:TUA5* reporter was introduced in *spo11-1* allowing us to follow the meiotic progression by using live cell imaging and MT state based determination of meiotic phases from a total of 224 observed meiocytes (**Supplemental Movie 5, Table 1, Supplemental Table 1**).

Interestingly and not previously recognized, *spo11-1* mutants spent much longer in early prophase (MT array state 2-3-4, late leptotene till early pachytene) compared to the wildtype (845 min, **Table 1, Figure 2A’**), with a predicted median time of 1119 min (CI 1031-1206 min, **Figure 6A**). The underlying molecular reason for this is not clear but interesting to study in the future. Important for this study, in *spo11-1* the duration of MT array state 5-6 had no statistical difference compared to the wildtype (360 min, **Table 1**, **Figure 2D′**), with a predicted median time of 374 min (CI 349-399 min, **Figure 6D**). This suggests that the complete loss of recombination caused by the absence of DSBs in the *spo11-1* mutant does not lead to the prolongation of MT array state 5-6 at 21°C.

**Figure 6.**
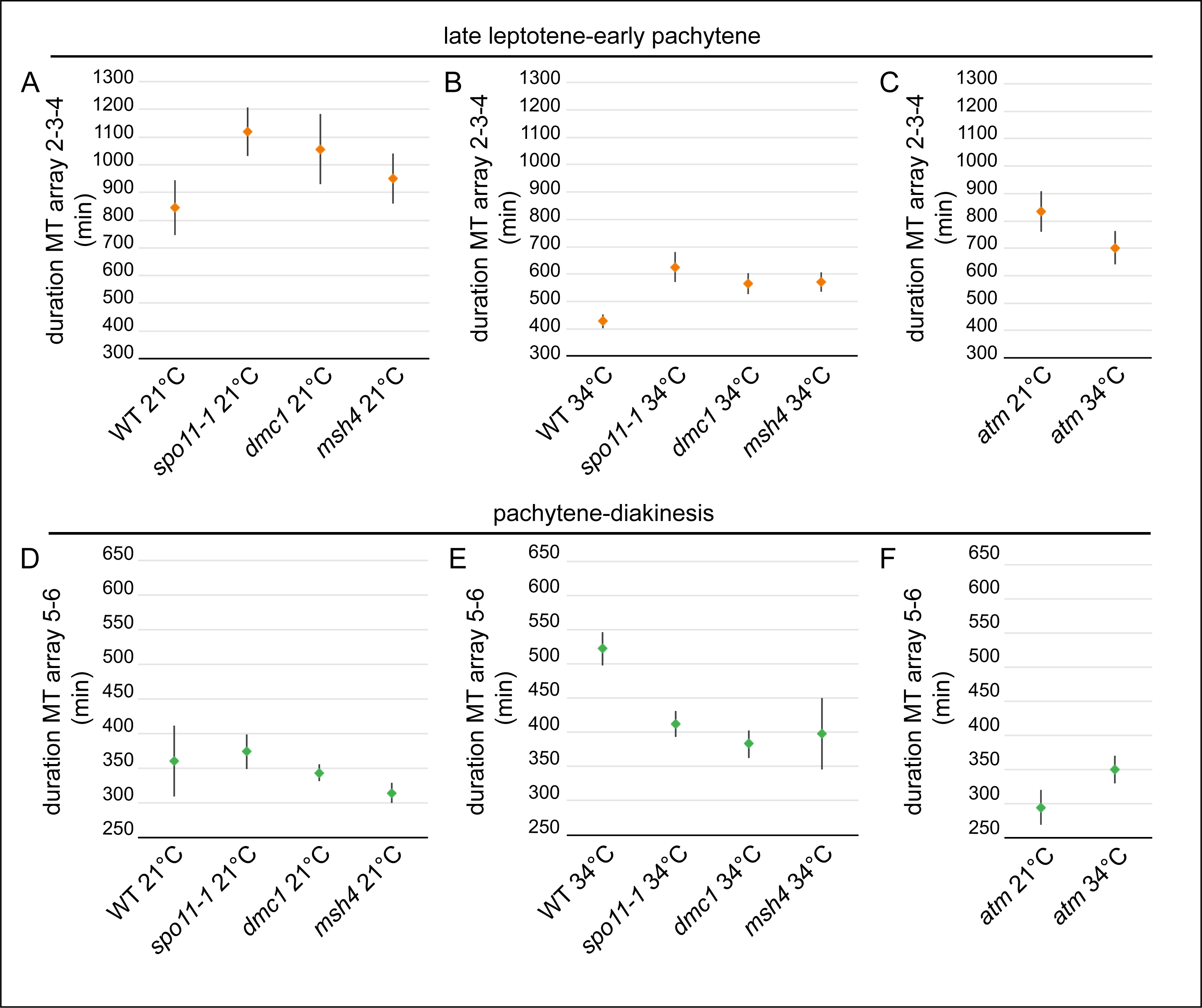
Duration of prophase in recombination mutants *spo11-1, dmc1, msh4* and *atm* at 21°C and HS34°C. The predicted median times (in min) with 95% confidence intervals of late leptotene-early pachytene (MT array state 2-3-4; orange; A-C) and pachytene-diakinesis (MT array state 5-6; green; D-F) in wildtype (as shown in Figure 2Á,B′) and recombination mutants *spo11-1, dmc1, msh4* at 21°C (A, D) and HS34°C (B, E) and the *atm* mutant at 21°C and HS34°C (C, F).

After prophase I, the meiotic division in *spo11-1* mutants continues with a predicted median time of 72 min (CI 67-76 min) for MT array state 7-8-9, followed by MT array state 10-11 for 63 min (CI 58-67 min), MT array state 12-13 for 48 min (CI 45-52 min) and finally MT array state 14 for 356 min (CI 326-385 min) (**Supplemental Figure 4**). Interestingly and unexpectedly, the durations of MT array state 7-8-9, MT array state 10-11 and MT array state 14 of *spo11-1* mutants show a statistical difference to the wildtype (**Figure 2Ć, D′, F′**).

Next, we asked whether a prolongation of MT state 5-6 could be dependent on homologous recombination repair by following meiosis in *dmc1* mutants in which we introduced the *TagRFP:TUA5* reporter and observed a total of 157 meiocytes (**Supplemental Movie 6, Table 1, Supplemental Table 1**). In *dmc1* mutants, DSBs are repaired through the sister chromatid of the same chromosome in an HR-dependent manner (Kurzbauer et al., 2012). The predicted median time per state was calculated and resulted in a duration of 1056 min for MT array state 2-3-4 (CI 929-1184 min, **Figure 6A**), showing a statistical difference to the wildtype (845 min, **Figure 2A’**) and resembling the extension of this phase seen in *spo11-1*. Thus, loss of early recombination steps appears to trigger a prolongation of early meiosis. For MT array state 5-6 in *dmc1* we determined a similar duration of 343 min (CI 331-355 min, **Figure 6D**) compared to wildtype (360 min, **Figure 2B′**), hence, for *dmc1* mutants we also do not observe a temporal extension of MT state 5-6. The meiotic division continued with a median time of 67 min (CI 63-71 min) for MT array state 7- 8-9. MT state 10-11 takes 63 min (CI 59-67 min), MT array state 12-13 lasts 47 min (CI 45-49 min) and MT array state 14 spans 281 min (CI 262-301 min, **Supplemental Figure 4**). All these subsequent phases are similar to its duration in *spo11-1,* which is interesting to investigate in the future.

Finally, we tested whether a failure to resolve recombination intermediates as Type I COs could be responsible for the delayed onset of NEB, using *msh4* mutants that contain the *TagRFP:TUA5* reporter (**Supplemental Movie 7**). A total of 193 meiocytes were and for every stage, the predicted median time was calculated (**Table 1, Supplemental Table 1**). In *msh4*, the MT array state 2-3-4 takes 951 min (CI 861-1040 min). This duration is not statistically different from wildtype (845min, **Figure 2A′**) but lies in between the CI for the wildtype on the one hand and the CI for *spo11-1* and *dmc1* on the other hand. Hence, it is difficult to judge from this dataset so far, whether this extension is relevant in comparison to the wildtype and resembles the situation found in the other two recombination mutants.

Subsequently, in *msh4,* a duration of 314 min (CI 299-329 min) was determined for MT array state 5-6 (**Figure 6A,D**). The meiotic division continued with extended MT array state 7-8-9 for 67 min (CI 65-72 min) compared to wildtype (47 min, **Figure 2C′**), which is similar as seen for *spo11-1* and *dmc1.* Next, MT array state 10-11 lasts 59 min (CI 56-63 min), MT array state 12-13 takes 49 min (CI 46-52 min) and finally, MT array state 14 spans 274 min (CI 253-294 min, **Supplemental Figure 3**). Thus, all recombination mutants tested have a similar duration of MT array states 7-8-9, 10-11, 12-13 and 14, compared to the wildtype (**Figure 2**). Yet, *msh4* mutants progressed through pachytene/diakinesis as wild-type plants at 21°C. Previously, 5‘-bromo-2‘-deoxyuridine (BrdU) labeling experiments showed a delay of S-phase till the end of prophase I in *msh4* mutants of 8 h, which could not be confirmed here, most probably because our time predictions did not include meiotic S-phase and early leptotene, where MSH4 is known to start appearing as numerous foci on the axes (Higgins et al., 2004).

Taken together, these results indicate that the loss of recombination *per se* does not cause the elongation of the MT array state 5-6 seen in wild-type meiocytes at 34°C.

### Prolongation of MT array state 5-6 at very high temperature is recombination-dependent

To further investigate the role of the recombination pathway on the elongation of MT array state 5-6 upon very high temperature heat stress, we observed a total of 198 meiocytes and analyzed the duration of this phase in *spo11-1* mutants at 34°C (**Supplemental Movie 8, Table 1, Supplemental Table 1**). Identically to the heat shock treatments of wild-type meiocytes, only flower buds which were in MT array state 1 were used for the modelling of the duration of the different meiotic states at HS34°C.

The MT array state 2-3-4 had a predicted median time of 626 min (CI 572-681 min), which is a decrease in duration compared to *spo11-1* at 21°C (1119 min, **Figure 6A**), showing a similar reduction as described for wild-type meiocytes (**Figure 6B**). Notably, the elongation of MT state 5-6 seen in the wildtype at HS34°C (522 min, a delay of 126 min compare to wildtype 21°C, **Figure 2B′**) was not found in *spo11-1* mutant at HS34°C, with a predicted median time of 412 min (CI 393-431 min, **Figure 6E**), compared to *spo11-1* at 21°C (374 min, **Figure 6D**). Further, MT array state 7-8-9 takes 35 min (CI 32-38 min), MT array state 10-11 spans 54 min (CI 49-58 min) and MT array state 12-13 lasts 23 min (CI 21-26 min, **Supplemental Figure 4**). All these states, except MT array state 5-6, had no statistical difference compared to wildtype at HS34°C (**Figure 2**), showing a similar reduction as described for wildtype at 21°C and HS34°C. For MT array state 14 of *spo11-1* at HS34°C, we could not provide the median time, identical to wildtype at HS34°C.

This suggested that the delay in the wildtype at very high temperature is not due to the absence of recombination but likely due to aberrant recombination intermediates and when they cannot be formed, as in *spo11-1* mutants, meiosis progresses without delay. To corroborate this, we next observed a total of 160 *dmc1* and 116 *msh4* meiocytes and measured the duration of MT state 5-6 at HS34°C (**Supplemental Movies 9-10, Table 1, Supplemental Table 1**). The predicted median time of MT array state 2-3-4 of *dmc1* upon HS34°C was 565 min (CI 526-605 min) and for the *msh4* mutant 571 min (CI 536-606 min, **Figure 6B**), which is a decrease in median time compared to these mutants at 21°C (**Figure 6A**). The predicted median time of MT array state 5-6 of *dmc1* and *msh4* at HS34°C was 383 min (CI 362-403 min) and 398 min (CI 346-450 min), respectively (**Figure 6E**). Compared to these mutants at 21°C, the duration was slightly longer, yet statistically different, but not to the degree of the elongation seen in wild-type meiocytes at 34°C (**Figure 2B′**).

These mutants at 34°C continued the meiotic division with MT array state 7-8-9 for 30 min (CI 28-32 min) and 32 min (CI 30-34 min), MT array state 10-11 for 57 min (CI 54-60 min) and 49 min (CI 43-55 min) and MT array state 12-13 for 22 min (CI 21-23 min) and 24 min (CI 22-26 min), respectively (**Supplemental Figure 3**). Similar to wildtype and *spo11-1* at HS34°C, we could not provide a predicted median time for the MT array state 14.

Taken together, the absence of the 2.7 h prolonged duration of MT array state 5-6 in the recombination mutants, *spo11-1*, *dmc1* and *msh4,* upon HS34°C leads to the hypothesis that the extension of the MT array state 5-6 upon 34°C is recombination dependent.

### High temperature reveals the presence of a pachytene checkpoint in Arabidopsis

The here observed prolongation of pachytene/diakinesis is reminiscent of the pachytene checkpoint in animals and yeast. However, the observation that mutants devoid of recombination progress through meiosis in plants, as quantified above, has previously raised the hypothesis that plants do not have a pachytene checkpoint (Caryl et al., 2003; Jackson et al., 2006; Li et al., 2009b). A central executer of the pachytene checkpoint in yeast and animals is the checkpoint kinase ATM (Lange et al., 2011; Pacheco et al., 2015; Penedos et al., 2015). ATM is highly conserved and also plays a major role in meiosis in Arabidopsis, for instance for the repair of DSBs (Garcia et al., 2003; Kurzbauer et al., 2021; Lange et al., 2011; Li et al., 2004; Yao et al., 2020).

To test an involvement of ATM in the prolongation of pachytene/diakinesis upon heat stress, we introduced the *TagRFP:TUA5* reporter in the *atm* mutant and followed meiotic progression at 21°C and HS34°C using live cell imaging and observed a total of 228 and 172 meiocytes, respectively, and determined the duration of the MT array states as described before (**Supplemental Movies 11-12, Table 1, Supplemental Table 1**). At 21°C, the MT array state 2-3-4 lasts 834 min (CI 761-908 min) and MT array state 5-6 takes 295 min (CI 270-321, **Figure 6C,F**). Thus, *atm* meiocytes progressed even faster than the wildtype through MT array state 5-6 (360 min, **Figure 2B′**), hinting at a possible role in prolonging this phase even under control conditions.

Next, MT array state 7-8-9 takes 45 min (CI 42-49 min), MT array state 10-11 lasts 60 min (CI 55-66 min), MT array state 12-13 persists 43 min (CI 40-46 min) and MT array state 14 spans 245 min (CI 230-260 min, **Supplemental Figure 3**). The biological relevance of the statistical difference of MT array state 10-11 and 14, compared to wildtype, needs to be further investigated.

Under HS34°C, the MT array state 2-3-4 had a predicted median time of 702 min (CI 640-764 min) and surprisingly the prolongation of MT array state 5-6 seen in the wildtype was abolished, *i.e.* 350 min (CI 330-370 min) in *atm* mutants versus 522 min in the wildtype (**Table 1**, **Figure 6C,F**). Thus, *atm* meiocytes progressed through this phase with a similar speed as meiocytes in which recombination is abolished.

All other MT array states were not found to be obviously different in duration when compared to the wildtype at 34°C, *i.e.* MT array state 7-8-9 lasts 31 min (CI 29-33 min), MT array state 10-11 takes 55 min (CI 50-60 min) and MT array state 12-13 spans 26 min (CI 23-28 min, **Figure 6C,F**, **Supplemental Figure 3**). Once again, the duration of MT array state 14 at HS34°C could not be predicted.

These results suggest the involvement of ATM in the prolongation of pachytene/diakinesis upon 34°C. Given the similarities in extension of pachytene/diakinesis and the involvement of ATM, we conclude that Arabidopsis and likely other plants do have a specialized variant of the pachytene checkpoint that relies on the action of ATM and possibly other regulators to monitor aberrant recombination intermediates at high temperatures but, in contrast to animals, not the absence of recombination.

## DISCUSSION

More than 50 years ago, the consequences of high temperature on plant development in general and on meiosis in particular were already studied (Dowrick, 1957; Pao and Li, 1948; Wilson, 1959). Due to the latest insights into climate change, research on the influence of temperature on meiosis has been revived. Previous and current studies have relied on the analysis of fixed samples and obtained important insights into the duration of meiosis and meiotic recombination patterns at elevated temperatures. Here, we have followed a complementary approach by following meiosis by time-lapse live cell imaging. This has allowed us to obtain a highly temporally resolved dissection of meiotic progression in which we have compared the effects of three heat stress treatments, *i.e.* a heat shock at 30°C and 34°C and a long-term (one week) treatment at 30°C in comparison to the control temperature of 21°C. Notably, this work has provided novel insights into the effects of temperature on recombination as well as meiotic progression and has set the stage for revising of a dogma in the field.

### Formation of stress granules in meiosis

Heat stress induces a multitude of cellular responses, including the inhibition of general translation and the formation of SGs, which are proposed to function as transient places for both storage and degradation of proteins and mRNAs during stress resulting in a re-programming of translation. This is thought to be especially important for the re-initiation of translation upon recovery from the stress condition, as reviewed by (Anderson and Kedersha, 2002, 2008; Buchan and Parker, 2009). In mice spermatocytes, SGs have been previously found to be formed after heat treatment (42°C) and these SGs contain for instance DAZL, an RNA-binding protein, which interacts with the SC, is involved in mRNA transport and is proposed to function as a translational activator (Kim et al., 2012).

Labelling the major cell cycle regulator of Arabidopsis, CDKA;1, we have shown here that meiocytes in Arabidopsis also form SGs at 30°C and 34°C. CDKA;1 has been previously demonstrated along with several other proteins, like MPK3 and TORC1, to be present in SGs of heat-stressed seedlings (Kosmacz et al., 2019). Why and how CDKA;1 is recruited to the SGs and which other proteins and RNAs are present in SGs during meiosis remains to be investigated. It has been previously hypothesized that the presence of CDKA;1 in SGs would allow a cell to resume cell division activity in Arabidopsis after attenuation of the stress (Kosmacz et al., 2019). CDKs typically require a co-factor, called cyclin, for their activity and in budding yeast, WHI8, an RNA-binding protein, was shown to bind to and recruit the mRNA of the cyclin CLN3 to SGs upon heat stress causing the inhibition of CLN3 translation (Yahya et al., 2021). Interestingly, CDC28, the homolog of CDKA;1 in budding yeast, itself is also recruited to SGs by WHI8 and has been found to play an important role in SG dissolution and the translation of SG-recruited mRNAs, such as the one of CLN3, upon release from stress.

This raises the question whether in Arabidopsis CDKA;1 is also a mediator of SG dissolution and subsequent re-initiation of translation. Interestingly, many proteins related to translation have been previously identified as putative CDKA;1 substrates (Pusch et al., 2011). A pivotal role of translational control for the abundance of proteins in meiosis has been established in several organisms including budding yeast (Brar et al., 2012). This gives rise to the speculation that translational regulation of meiosis in Arabidopsis is also present and likely controlled by CDK activity.

The appearance of CDKA;1 in SGs has allowed us to faithfully confirm the application of the heat stress in meiocytes. On the one hand, we could show that the heat stress reaches meiocytes relatively fast, *i.e.* in less than 15 min. Thus, our imaging starts when meiocytes are already exposed to the desired temperature applied in our set-up. On the other hand, we observed that SGs are not regularly found at 30°C. Thus, the appearance of SGs is also a sensor of the applied temperature itself. Since SGs are formed rapidly at 34°C, we assumed that the heat stress at 30°C also reaches meiocytes in a similar time frame providing confidence that we have been looking at an immediate effect of the high temperature rather than a ramping effect over a long period. We anticipate that the formation of CDKA;1-containing SGs could be used as a general readout to study heat stress in other plant tissues and possibly other plant species as well.

Interestingly, the localization of CDKA;1 to SGs is stage-specific and its SG localization could only be observed from pachytene onwards but not earlier in meiosis. Notably, DAZL also shows a stage-specific localization to SGs in mice spermatocytes and is recruited to SGs only during pachytene in response to heat, coinciding with its highest accumulation level (Kim et al., 2012). In comparison, CDKA;1 is dynamically localized in the nucleus and the cytoplasm and the formation of CDKA;1-positive SGs appears when its cytoplasmic portion is the highest. Whether the formation of CDKA;1-positive SGs is a hence of function of its cytoplasmic concentration or whether this relies on other stage specific parameters needs to be determined. Conversely, it is also not clear whether non-CDKA;1-containing SGs are formed prior to pachytene.

### Heat and meiotic progression

The durational changes of meiosis upon high temperatures were studied in several plant species including Arabidopsis, barley, wheat, *Dasypyrum villosum (L.) P. candargy* and bluebell (Bennett et al., 1972; Draeger and Moore, 2017; Higgins et al., 2012; Stefani and Colonna, 1996; Wilson, 1959). These studies have relied on static analysis of fixed material, *e.g*. anther fixation and staging before and after a certain time interval or BrdU pulse labelling followed by the analysis of meiotic chromosome figures (Armstrong et al., 2003; Bennett et al., 1972). These studies concluded that the duration of meiosis at high temperatures is decreased. Here, we have confirmed the general trend of increased speed of meiosis at high temperatures. However, our live cell imaging approach allowed us to follow meiotic progression with unprecedented depth generating quantitative data that can be statistically analyzed. This led to the finding not all meiotic phases respond equally to an increase in temperature. Most strikingly, we found that pachytene/diakinesis were substantially extended at 34°C when compared to control conditions at 21°C, as seen by a considerable prolongation of the time of NEB.

This opens the door to study which regulators and/or processes are sensitive to heat. For instance, it is well-established that NEB in animals is under full control of CDK activity and nuclear envelope components, like lamins are bona fide CDK substrates (Adhikari et al., 2012; Gong et al., 2007; Zuela and Gruenbaum, 2016). How NEB is controlled in plants is still an enigma, especially also since lamins do not appear to be conserved in plants (Ciska and Moreno Diaz de la Espina, 2013; Fiserova and Goldberg, 2010). However, it is tempting to speculate that NEB is also under the control of CDK activity. Hence, the here-observed delay in NEB might be directly or indirectly mediated by a repression of CDK activity.

NEB also likely represents a gate in meiotic progression. Chromosomes are strong microtubule organizing structures in plants (Lee and Liu, 2019), and once the nuclear envelope is broken down, the MT array that is enriched around the nucleus quickly connects to the chromosomes and organizes itself into a spindle (Prusicki et al., 2019). Thus, a delay of NEB represents a physical barrier that provides additional time to complete and/or correct processes in the reaction environment of the nucleus before chromosomes start to be moved in the cell.

### Heat and meiotic recombination

To further explore the extension of pachytene/diakinesis under heat stress, we have genetically and temporally dissected this effect. An obvious cause for the observed prolongation was altered meiotic recombination, supported by our study and previous analyses of meiotic chromosome configurations (De Storme and Geelen, 2020; Hedhly et al., 2020; Higgins et al., 2012). Using then mutants of genes that control different steps in the meiotic recombination process, like *spo11-1*, *dmc1*, and *msh4*, we have shown that the extension of pachytene/diakinesis is recombination dependent, *i.e*. the extension of pachytene/diakinesis was lost in these mutants at 34°C. Notably, these mutants, when grown under non-stress conditions at 21°C, do not display a relevant prolongation of meiosis in a way that we could detect with our assays. This stands in contrast to animals where loss of recombination, *e.g*. in *dmc1* mice mutants, triggers meiotic arrest and subsequently induces cell death (Barchi et al., 2005; de Rooij and de Boer, 2003; Roeder and Bailis, 2000).

To further narrow down the origin of the elongation of pachytene/diakinesis, we applied heat stress only around zygotene, *i.e.* up to 17 h later than in our first sets of experiments. Notably, this late heat stress did not cause a prolongation and hence, recombination appears to be affected prior to SC formation. This is interesting since earlier work indicated that the SC is severely affected by heat leading to so-called polycomplexes in which transverse filaments become laterally connected and a study in *C. elegans* suggests that ZYP1 aggregation upon high temperature primarily reflects SC assembly failure (Bilgir et al., 2013; Higgins et al., 2012; Loidl, 1989). In addition, temporal dissection of heat stress on grasshopper spermatocytes revealed that heat-induced chiasma frequency changes are most likely the consequence of the completeness or efficiency of pairing (Henderson, 1988). Thus, we conclude that already very early recombination processes, such as pairing of homologs, are affected by heat and that likely these aberrant processes cause the formation of polycomplexes. However, it is still possible that aberrant SC configuration ZYP1 can serve as a signal to cause a delay in NEB.

From our mutant analysis and chromosome spreads at elevated temperatures, it is likely that recombination intermediates cause this delay. What the structure of these intermediates is and how they cause a delay needs to be investigated in the future. It is possible that the delay is triggered by non-homologous recombination caused by mispairing and hence partially interconnected chromosomes. A *zmm* mutant analysis in yeast revealed that a specific block in progression of CO formation occurs at high temperatures, resulting in the formation of intermediates and/or interactions with sister chromatids (Borner et al., 2004). Further, it is well known from yeast that unresolved recombination intermediates can cause nuclear division defects (Kaur et al., 2015; Kaur et al., 2019; Tang et al., 2015).

At high temperature (reported up to 33°C), DSB formation remains unaffected in yeast and Arabidopsis, whereas from *C. elegans* spermatocytes it is known that high temperature (above or at the threshold of 34°C) induces SPO11-1-independent DSBs, which are recognized by the CO repair machinery (Brown et al., 2020; Kurhanewicz et al., 2020; Modliszewski et al., 2018). Whether Arabidopsis has a different recombination effect/response below and above a temperature threshold and if there is a different molecular mechanism remains to be investigated in the future.

### A specialized pachytene checkpoint in Arabidopsis

Aberrant recombination structures and the absence of recombination trigger meiotic arrest in animals and yeast, this arrest is controlled by the so-called pachytene checkpoint causing meiotic arrest in early pachytene (Barchi et al., 2005; Bishop et al., 1992; Rockmill et al., 1995). Since in plants mutants in which recombination is abolished, such as *dmc1*, are not arrested in meiosis, it was proposed that plants do not have a pachytene checkpoint (Couteau et al., 1999; Grelon et al., 2001; Higgins et al., 2004; Li et al., 2004).

A major regulator of the pachytene checkpoint in animals and yeast is the checkpoint kinase ATM (Barchi et al., 2005; Lange et al., 2011; Pacheco et al., 2015; Penedos et al., 2015; Roeder and Bailis, 2000). Removing ATM in mutants that trigger the pachytene checkpoint, for instance in weak loss-of-function mutants for Trip13/PCH2, promotes further progression through pachytene indicating that the early arrest is under control of this checkpoint kinase (Pacheco et al., 2015).

In budding yeast, *atm* mutants undergo the first meiotic division before all recombination events are complete (Lydall et al., 1996; Stuart and Wittenberg, 1998). Indeed, we found that the pachytene/diakinesis extension is lost in *atm* mutants implicating ATM in this checkpoint and the execution of the observed meiotic delay, *e.g*. by sensing of aberrant recombination structures. Taken together with our finding that the prolongation of pachytene/diakinesis is recombination dependent, we conclude that Arabidopsis and likely other plants do have a pachytene checkpoint. However, this checkpoint is less stringent than in animals since it does not respond to the absence of meiotic recombination. Moreover, the extension is timely restricted and typically after 2.7 h meiosis continues. It needs to be analyzed in the future, when the nature of the presumptive aberrant recombination intermediates is understood, whether they are resolved during this time or whether the checkpoint erodes, *i.e.* even though checkpoint conditions are not fulfilled, meiosis progresses. An erosion has been found for another checkpoint in plants, *i.e*. the spindle assembly checkpoint that assures that all chromosomes are connected to microtubule fibers of the spindle. Triggering this checkpoint was only able to delay anaphase onset by a maximum of less than 2 h (Komaki and Schnittger, 2017).

It is an interesting discussion point whether less stringent cell division checkpoints (pachytene and spindle assembly checkpoint (SAC)) represent an evolutionary strategy in plants. Genome mutations, especially polyploidization events are more prominent in plants than in animals and are suspected to be a major driving force of evolution in plants (Brownfield and Kohler, 2011; De Storme and Geelen, 2013; Li et al., 2009b; Wijnker and Schnittger, 2013). Moreover, hybridization events are very frequent in plants. An alien genome would likely affect recombination by either reducing it or causing aberrant recombination structures. Less stringent checkpoints would pave the road for hybridization events since by chance viable combinations of chromosomes are generated. Especially an interplay between a relaxed pachytene checkpoint and a relaxed SAC would promote rapid genome evolution.

## MATERIAL AND METHODS

### Plant material and growth conditions

All *Arabidopsis thaliana* plants used in this study were derived from the Columbia (Col-0) ecotype. The *CDKA;1:mVenus–TagRFP:TUA5* double reporter line, *KINGBIRD reporter line 2* (*PRO_REC8_:REC8:GFP/PRO_RPS5A_:TagRFP:TUA5*) and the *ASY1:RFP-ZYP1b:GFP* double reporter line have been previously described (Prusicki et al., 2019; Sofroni et al., 2020; Yang et al., 2019; Yang et al., 2020). The T-DNA insertion lines for *DMC1* (GABI_918E07), *SPO11-1* (SALK_146172), *MSH4* (SALK_136296) and *ATM* (SALK_006953) were obtained from GABI-Kat T-DNA mutation collection via NASC (http://arabidopsis.info/) and the collection of T-DNA mutants at the Salk Institute Genomic Analysis Laboratory (http://signal.salk.edu/cgi-bin/tdnaexpress).

Seeds were surface-sterilized with chlorine gas and germinated on 1% agar containing half-strength Murashige and Skoog (MS) salts and 1% sucrose, pH 5.8. When required, antibiotics were added for seed selection. All plants were grown under long-day conditions (16 h light at 21°C (+/- 0.5 °C)/ 8 h dark at 18°C (+/- 0.5 °C), with 60% humidity). For short-term heat treatment, plants were first grown under standard long-day conditions until sample preparations for live cell imaging or transferred to a climate chamber at long-day photoperiod with continuous temperature (30°C/34°C (+/- 0.5°C)) for 16/24 h prior to observation/fixation. For long-term heat treatment, healthy plants at bolting stage were transferred to a climate chamber at long-day photoperiod with a continuous temperature of 30°C (+/- 0.5°C) with 60% humidity for 7 days.

### Plasmids and plant transformation

The reporter constructs *PRO_RPS5A_:TagRFP:TUA5* and *KINGBIRD reporter line 2*, previously described (Prusicki et al., 2019), were transformed into the T-DNA insertion plants by floral dipping and T1 seeds were selected on half strength MS supplemented with antibiotics hygromycin, from T2 generation onwards plants were used for observation.

### Confocal microscopy and intensity plot

For protein localization experiments, healthy flower buds were dissected exposing 2 anthers and carefully positioned in a petri dish, with 0.8% agar with half-strength MS salts, pH 5.8, and meiocytes of different meiotic stages were imaged using a Zeiss LSM880 confocal microscope.

For the pixel intensity plot, flower buds were dissected and the anthers in MT array state 6 were imaged using a Zeiss LSM880 confocal microscope with the exact same settings for the different heat conditions. The pixel brightness was measured through a region of interest using ImageJ and plotted against the X dimension, which is the distance of the region of interest.

### Live cell imaging and data processing

Live cell imaging was performed as described previously (Prusicki et al, 2019). In short, up to 6 flower buds of 0.2-0.6 mm were carefully positioned in a petri dish with 0.8% agar with half-strength MS salts, pH 5.8. Time lapse was performed using an upright Zeiss LSM 880 confocal microscope with ZEN 2.3 SP1 software (Carl Zeiss AG, Oberkochen, Germany) and a W-plan Apochromat 40X/ 1.0 DIC objective (Carl Zeiss AG, Oberkochen, Germany). GFP and TagRPF were excited at λ= 488 nm and 561 nm, respectively, and detected between 498-560 nm and 520-650 nm, respectively. Auto-fluorescence was detected between 680-750 nm. With a time interval of 10 min, a series of 6 Z-stacks with 50 μm distance was acquired under a thermally controlled environment (21°C/30°C/34°C (+/- 0.15%)) in an incubation chamber. Due to sample movement, the Z-planes were manually selected using the review multi-dimensional data function of the software Metamorph Version 7.8 and the XY movement was corrected using the Stack Reg plugin of Fiji.

### Quantitative analysis of the meiotic phases

The analysis of the duration is based on the *TagRFP:TUA5* reporter. Meiocytes were manually assigned to defined MT states. The data were collected from a minimum of 3 independent set-ups, with a minimum of 8 anthers per genotype per heat treatment. The durations of the meiotic phases were extracted from at least 65 meiocytes.

### Statistical methods

The chosen distributions underlying our parametric model were log-normal for MT array states 7-8-9, 10-11, 12-13 and 14, whereas for MT array states 2-3-4 and 5-6 and the model for the complete duration of MT array states 2-13 a Weibull distribution was selected. Estimation results are presented as predicted marginal median times, together with 95% confidence intervals. The statistical analysis was performed with R version 3.5.1 and Stata SE version 16.1.

### Cytology

The cytological analysis of the meiocytes under short and long heat treatment was done by performing chromosome spreads, as previously described (Sofroni et al., 2020). Briefly, healthy flower buds were fixated in 3:1 ethanol:acetic acid for a minimum of 24 h at 4°C, following washing steps with 70% ethanol and stored at 4°C. Next, flower buds were washed in water and in 10 mM citrate buffer, pH 4.5 and digested in an enzyme mix (10 mM citrate buffer containing 0.5% cellulase, 0.5% pectolyase and 0.5% cytohelicase) for 2.5 h at 37°C. Digested flower buds were squashed and spread onto a glass slide in 45% acetic acid on a 46°C hotplate. Finally, the slides were washed in cold 3:1 ethanol:acetic acid and mounted in Vectashield medium with DAPI (Vector Laboratories).

### Supplemental Data files

Supplemental Table 1 Detailed overview of the sample size

Supplemental Figure 1 Microtubule array states upon heat stress

Supplemental Figure 2 Meiotic defects of wildtype upon heat stress

Supplemental Figure 3 Duration of meiotic phases in recombination mutants *spo11-1*, *dmc1*, *msh4* and *atm* at 21°C and HS34°C

Supplemental Movie 1- The meiotic division of wild-type meiocytes at 21°C

Supplemental Movie 2- The meiotic division of wild-type meiocytes at HS30°C

Supplemental Movie 3- The meiotic division of wild-type meiocytes at HS34°C

Supplemental Movie 4- The meiotic division of wild-type meiocytes at LT30°C

Supplemental Movie 5- The meiotic division of *spo11-1* meiocytes at 21°C

Supplemental Movie 6- The meiotic division of *dmc1* meiocytes at 21°C

Supplemental Movie 7- The meiotic division of *msh4* meiocytes at 21°C

Supplemental Movie 8- The meiotic division of *spo11-1* meiocytes at HS34°C

Supplemental Movie 9- The meiotic division of *dmc1* meiocytes at HS34°C

Supplemental Movie 10- The meiotic division of *msh4* meiocytes at HS34°C

Supplemental Movie 11- The meiotic division of *atm* meiocytes at 21°C

Supplemental Movie 12- The meiotic division of *atm* meiocytes at HS34°C

## AUTHOR CONTRIBUTIONS

J.D.J.-B. and A.S. designed the research; J.D.J.-B. performed the experiments; L.K. and A.B. performed the statistical analysis; J.D.J.-B., L.K., A.B. and A.S. analyzed and discussed the data; J.D.J.-B. and A.S. wrote the article; J.D.J.-B., L.K., A.B. and A.S. revised and approved the article.

## AKNOWLEDGEMENTS

This research was funded by the University of Hamburg. We thank Lucas Lang (University of Hamburg) and Konstantinos Lampou (University of Hamburg) for critical reading and helpful comments on the manuscript. We are grateful to Chao Yang, Shinichiro Komaki and Konstantinos Lampou for providing the reporter lines in the mutant background.

## Parsed Citations

Adhikari, D., Zheng, W., Shen, Y., Gorre, N., Ning, Y., Halet, G., Kaldis, P., and Liu, K. (2012). Cdk1, but not Cdk2, is the sole Cdk that is essential and sufficient to drive resumption of meiosis in mouse oocytes. Hum Mol Genet 21, 2476–2484.

Anderson, P., and Kedersha, N. (2002). Visibly stressed: the role of eIF2, TIA-1, and stress granules in protein translation. Cell Stress Chaperones 7, 213–221.

Anderson, P., and Kedersha, N. (2008). Stress granules: the Tao of RNAtriage. Trends Biochem Sci 33, 141–150.

Anderson, T.R., Hawkins, E., and Jones, P.D. (2016). CO2, the greenhouse effect and global warming: fromthe pioneering work of Arrhenius and Callendar to today’s Earth System Models. Endeavour 40, 178–187.

Armstrong, S.J., Caryl, A.P., Jones, G.H., and Franklin, F.C. (2002). Asy1, a protein required for meiotic chromosome synapsis, localizes to axis-associated chromatin in Arabidopsis and Brassica. J Cell Sci 115, 3645–3655.

Armstrong, S.J., Franklin, F.C.H., and Jones, G.H. (2003). A meiotic time-course for Arabidopsis thaliana. Sexual Plant Reproduction 16, 141–149.

Bannigan, A., Scheible, W.R., Lukowitz, W., Fagerstrom, C., Wadsworth, P., Somerville, C., and Baskin, T.I. (2007). Aconserved role for kinesin-5 in plant mitosis. J Cell Sci 120, 2819–2827.

Barchi, M., Mahadevaiah, S., Di Giacomo, M., Baudat, F., de Rooij, D.G., Burgoyne, P.S., Jasin, M., and Keeney, S. (2005). Surveillance of different recombination defects in mouse spermatocytes yields distinct responses despite elimination at an identical developmental stage. Mol Cell Biol 25, 7203–7215.

Bennett, M.D., Smith, J.B., and Kemble, R. (1972). THE EFFECT OF TEMPERATURE ON MEIOSIS AND POLLEN DEVELOPMENT INWHEAT AND RYE. Canadian Journal of Genetics and Cytology 14, 615–624.

Berchowitz, L.E., Francis, K.E., Bey, A.L., and Copenhaver, G.P. (2007). The role of AtMUS81 in interference-insensitive crossovers in A. thaliana. PLoS Genet 3, e132.

Bilgir, C., Dombecki, C.R., Chen, P.F., Villeneuve, A.M., and Nabeshima, K. (2013). Assembly of the Synaptonemal Complex Is a Highly Temperature-Sensitive Process That Is Supported by PGL-1 During Caenorhabditis elegans Meiosis. G3 (Bethesda) 3, 585–595.

Bishop, D.K., Park, D., Xu, L., and Kleckner, N. (1992). DMC1: a meiosis-specific yeast homolog of E. coli recArequired for recombination, synaptonemal complex formation, and cell cycle progression. Cell 69, 439–456.

Bomblies, K., Higgins, J.D., and Yant, L. (2015). Meiosis evolves: adaptation to external and internal environments. New Phytol 208, 306–323.

Borner, G.V., Kleckner, N., and Hunter, N. (2004). Crossover/noncrossover differentiation, synaptonemal complex formation, and regulatory surveillance at the leptotene/zygotene transition of meiosis. Cell 117, 29–45.

Brar, G.A., Yassour, M., Friedman, N., Regev, A., Ingolia, N.T., and Weissman, J.S. (2012). High-resolution view of the yeast meiotic programrevealed by ribosome profiling. Science 335, 552–557.

Brown, S.D., Audoynaud, C., and Lorenz, A. (2020). Intragenic meiotic recombination in Schizosaccharomyces pombe is sensitive to environmental temperature changes. Chromosome Res 28, 195–207.

Brownfield, L., and Kohler, C. (2011). Unreduced gamete formation in plants: mechanisms and prospects. J Exp Bot 62, 1659–1668.

Buchan, J.R., and Parker, R. (2009). Eukaryotic stress granules: the ins and outs of translation. Mol Cell 36, 932–941.

Bulankova, P., Riehs-Kearnan, N., Nowack, M.K., Schnittger, A., and Riha, K.(2010). Meiotic progression in Arabidopsis is governed by complex regulatory interactions between SMG7, TDM1, and the meiosis I-specific cyclin TAM. Plant Cell 22, 3791–3803.

Cai, X., Dong, F., Edelmann, R.E., and Makaroff, C.A. (2003). The Arabidopsis SYN1 cohesin protein is required for sister chromatid arm cohesion and homologous chromosome pairing. J Cell Sci 116, 2999–3007.

Capilla-Perez, L., Durand, S., Hurel, A., Lian, Q., Chambon, A., Taochy, C., Solier, V., Grelon, M., and Mercier, R. (2021). The synaptonemal complex imposes crossover interference and heterochiasmy in Arabidopsis. Proc Natl Acad Sci U S A 118.

Caryl, A.P., Armstrong, S.J., Jones, G.H., and Franklin, F.C. (2000). Ahomologue of the yeast HOP1 gene is inactivated in the Arabidopsis meiotic mutant asy1. Chromosoma 109, 62–71.

Caryl, A.P., Jones, G.H., and Franklin, F.C. (2003). Dissecting plant meiosis using Arabidopsis thaliana mutants. J Exp Bot 54, 25–38.

Chodasiewicz, M., Sokolowska, E.M., Nelson-Dittrich, A.C., Masiuk, A., Beltran, J.C.M., Nelson, A.D.L., and Skirycz, A. (2020). Identification and Characterization of the Heat-Induced Plastidial Stress Granules Reveal New Insight Into Arabidopsis Stress Response. Front Plant Sci 11, 595792.

Ciska, M., and Moreno Diaz de la Espina, S. (2013). NMCP/LINC proteins: putative lamin analogs in plants Plant Signal Behav 8, e26669.

Collins, M. (2014). Long-term Climate Change: Projections, Commitments and Irreversibility Pages 1029 to 1076. In Climate Change 2013 – The Physical Science Basis: Working Group I Contribution to the Fifth Assessment Report of the Intergovernmental Panel on Climate Change, C. Intergovernmental Panel on Climate, ed. (Cambridge: Cambridge University Press), pp. 1029–1136.

Couteau, F., Belzile, F., Horlow, C., Grandjean, O., Vezon, D., and Doutriaux, M.P. (1999). Randomchromosome segregation without meiotic arrest in both male and female meiocytes of a dmc1 mutant of Arabidopsis. Plant Cell 11, 1623–1634.

de Rooij, D.G., and de Boer, P. (2003). Specific arrests of spermatogenesis in genetically modified and mutant mice. Cytogenet Genome Res 103, 267–276.

De Storme, N., and Geelen, D. (2013). Sexual polyploidization in plants--cytological mechanisms and molecular regulation. New Phytol 198, 670–684.

De Storme, N., and Geelen, D. (2020). High temperatures alter cross-over distribution and induce male meiotic restitution in Arabidopsis thaliana. Commun Biol 3, 187.

Dissmeyer, N., Nowack, M.K., Pusch, S., Stals, H., Inze, D., Grini, P.E., and Schnittger, A. (2007). T-loop phosphorylation of Arabidopsis CDKA;1 is required for its function and can be partially substituted by an aspartate residue. Plant Cell 19, 972–985.

Dowrick, G.J. (1957). The influence of temperature on meiosis. Heredity 11, 37–49.

Draeger, T., and Moore, G. (2017). Short periods of high temperature during meiosis prevent normal meiotic progression and reduce grain number in hexaploid wheat (Triticumaestivum L.). Theor Appl Genet 130, 1785–1800.

Dubiel, M., De Coninck, T., Osterne, V.J.S., Verbeke, I., Van Damme, D., Smagghe, G., and Van Damme, E.J.M. (2020). The ArathEULS3 Lectin Ends up in Stress Granules and Can Follow an Unconventional Route for Secretion. Int J Mol Sci 21.

Fiserova, J., and Goldberg, Martin W. (2010). Relationships at the nuclear envelope: lamins and nuclear pore complexes in animals and plants. Biochemical Society Transactions 38, 829–831.

France, M.G., Enderle, J., Rohrig, S., Puchta, H., Franklin, F.C.H., and Higgins, J.D. (2021). ZYP1 is required for obligate cross-over formation and cross-over interference in Arabidopsis. Proc Natl Acad Sci U S A 118.

Garcia, V., Bruchet, H., Camescasse, D., Granier, F., Bouchez, D., and Tissier, A. (2003). AtATM is essential for meiosis and the somatic response to DNAdamage in plants. Plant Cell 15, 119–132.

Gong, D., Pomerening, J.R., Myers, J.W., Gustavsson, C., Jones, J.T., Hahn, A.T., Meyer, T., and Ferrell, J.E., Jr. (2007). Cyclin A2 regulates nuclear-envelope breakdown and the nuclear accumulation of cyclin B1. Curr Biol 17, 85–91.

Grelon, M., Vezon, D., Gendrot, G., and Pelletier, G. (2001). AtSPO11-1 is necessary for efficient meiotic recombination in plants. EMBO J 20, 589–600.

Hamada, T., Yako, M., Minegishi, M., Sato, M., Kamei, Y., Yanagawa, Y., Toyooka, K., Watanabe, Y., and Hara-Nishimura, I. (2018). Stress granule formation is induced by a threshold temperature rather than a temperature difference in Arabidopsis. J Cell Sci 131.

Hartung, F., Wurz-Wildersinn, R., Fuchs, J., Schubert, I., Suer, S., and Puchta, H. (2007). The catalytically active tyrosine residues of both SPO11-1 and SPO11-2 are required for meiotic double-strand break induction in Arabidopsis. Plant Cell 19, 3090–3099.

Hatfield, J.L., and Prueger, J.H. (2015). Temperature extremes: Effect on plant growth and development. Weather and Climate Extremes 10, 4–10.

Hedhly, A., Nestorova, A., Herrmann, A., and Grossniklaus, U. (2020). Acute heat stress during stamen development affects both the germline and sporophytic lineages in Arabidopsis thaliana (L.) Heynh. Environmental and Experimental Botany 173, 103992.

Henderson, S.A. (1988). Four effects of elevated temperature on chiasma formation in the locust Schistocerca gregaria. Heredity 60, 387–401.

Higgins, J.D., Armstrong, S.J., Franklin, F.C., and Jones, G.H. (2004). The Arabidopsis MutS homolog AtMSH4 functions at an early step in recombination: evidence for two classes of recombination in Arabidopsis. Genes Dev 18, 2557–2570.

Higgins, J.D., Perry, R.M., Barakate, A., Ramsay, L., Waugh, R., Halpin, C., Armstrong, S.J., and Franklin, F.C. (2012). Spatiotemporal asymmetry of the meiotic programunderlies the predominantly distal distribution of meiotic crossovers in barley. Plant Cell 24, 4096–4109.

Higgins, J.D., Sanchez-Moran, E., Armstrong, S.J., Jones, G.H., and Franklin, F.C. (2005). The Arabidopsis synaptonemal complex protein ZYP1 is required for chromosome synapsis and normal fidelity of crossing over. Genes Dev 19, 2488–2500.

Interthal, H., and Heyer, W.D. (2000). MUS81 encodes a novel helix-hairpin-helix protein involved in the response to UV- and methylation-induced DNAdamage in Saccharomyces cerevisiae. Mol Gen Genet 263, 812–827.

Jachymczyk, W.J., von Borstel, R.C., Mowat, M.R., and Hastings, P.J.(1981). Repair of interstrand cross-links in DNAof Saccharomyces cerevisiae requires two systems for DNArepair: the RAD3 system and the RAD51 system. Mol Gen Genet 182, 196–205.

Jackson, N., Sanchez-Moran, E., Buckling, E., Armstrong, S.J., Jones, G.H., and Franklin, F.C. (2006). Reduced meiotic crossovers and delayed prophase I progression in AtMLH3-deficient Arabidopsis. EMBO J 25, 1315–1323.

Jones, G.H., and Franklin, F.C.H. (2008). Meiosis in Arabidopis thaliana: Recombination,Chromosome Organization and Meiotic Progression. In Recombination and Meiosis: Crossing-Over and Disjunction, R. Egel, and D.-H. Lankenau, eds. (Berlin, Heidelberg: Springer Berlin Heidelberg), pp. 279–306.

Kaur, H., De Muyt, A., and Lichten, M.(2015). Top3-Rmi1 DNAsingle-strand decatenase is integral to the formation and resolution of meiotic recombination intermediates. Mol Cell 57, 583–594.

Kaur, H., Gn, K., and Lichten, M. (2019). Unresolved Recombination Intermediates Cause a RAD9-Dependent Cell Cycle Arrest in Saccharomyces cerevisiae. Genetics 213, 805–818.

Keeney, S., Giroux, C.N., and Kleckner, N. (1997). Meiosis-specific DNAdouble-strand breaks are catalyzed by Spo11, a member of a widely conserved protein family. Cell 88, 375–384.

Kim, B., Cooke, H.J., and Rhee, K. (2012). DAZL is essential for stress granule formation implicated in germcell survival upon heat stress. Development 139, 568–578.

Komaki, S., and Schnittger, A. (2017). The Spindle Assembly Checkpoint in Arabidopsis Is Rapidly Shut Off during Severe Stress. Dev Cell 43, 172–185 e175.

Kosmacz, M., Gorka, M., Schmidt, S., Luzarowski, M., Moreno, J.C., Szlachetko, J., Leniak, E., Sokolowska, E.M., Sofroni, K., Schnittger, A., et al. (2019). Protein and metabolite composition of Arabidopsis stress granules. New Phytol 222, 1420–1433.

Kukal, M.S., and Irmak, S. (2018). Climate-Driven Crop Yield and Yield Variability and Climate Change Impacts on the U.S. Great Plains Agricultural Production. Sci Rep 8, 3450.

Kurhanewicz, N.A., Dinwiddie, D., Bush, Z.D., and Libuda, D.E. (2020). Elevated Temperatures Cause Transposon-Associated DNA Damage in C. elegans Spermatocytes. Curr Biol 30, 5007–5017 e5004.

Kurzbauer, M.T., Janisiw, M.P., Paulin, L.F., Prusen Mota, I., Tomanov, K., Krsicka, O., Haeseler, A.V., Schubert, V., and Schlogelhofer, P. (2021). ATM controls meiotic DNAdouble-strand break formation and recombination and affects synaptonemal complex organization in plants. Plant Cell.

Kurzbauer, M.T., Uanschou, C., Chen, D., and Schlogelhofer, P. (2012). The recombinases DMC1 and RAD51 are functionally and spatially separated during meiosis in Arabidopsis. Plant Cell 24, 2058–2070.

Lamb, B.C. (1969). Related and unrelated changes in conversion and recombination frequencies with temperature in Sordaria fimicola, and their relevance to hybrid-DNA models of recombination. Genetics 62, 67–78.

Lange, J., Pan, J., Cole, F., Thelen, M.P., Jasin, M., and Keeney, S. (2011). ATM controls meiotic double-strand-break formation. Nature 479, 237–240.

Lee, Y.J., and Liu, B. (2019). Microtubule nucleation for the assembly of acentrosomal microtubule arrays in plant cells. New Phytol 222, 1705–1718.

Lei, X., Ning, Y., Eid Elesawi, I., Yang, K., Chen, C., Wang, C., and Liu, B. (2020). Heat stress interferes with chromosome segregation and cytokinesis during male meiosis in Arabidopsis thaliana. Plant Signal Behav 15, 1746985.

Li, H., Zeng, X., Liu, Z.Q., Meng, Q.T., Yuan, M., and Mao, T.L. (2009a). Arabidopsis microtubule-associated protein AtMAP65-2 acts as a microtubule stabilizer. Plant Mol Biol 69, 313–324.

Li, W., Chen, C., Markmann-Mulisch, U., Timofejeva, L., Schmelzer, E., Ma, H., and Reiss, B. (2004). The Arabidopsis AtRAD51 gene is dispensable for vegetative development but required for meiosis. Proc Natl Acad Sci U S A 101, 10596–10601.

Li, X.C., Barringer, B.C., and Barbash, D.A. (2009b). The pachytene checkpoint and its relationship to evolutionary patterns of polyploidization and hybrid sterility. Heredity (Edinb) 102, 24–30.

Liu, B., De Storme, N., and Geelen, D. (2017). Cold interferes with male meiotic cytokinesis in Arabidopsis thaliana independently of the AHK2/3-AHP2/3/5 cytokinin signaling module. Cell Biol Int 41, 879–889.

Lloyd, A., Morgan, C., FC, H.F., and Bomblies, K. (2018). Plasticity of Meiotic Recombination Rates in Response to Temperature in Arabidopsis. Genetics 208, 1409–1420.

Loidl, J. (1989). Effects of elevated temperature on meiotic chromosome synapsis in Allium ursinum. Chromosoma 97, 449–458.

Lydall, D., Nikolsky, Y., Bishop, D.K., and Weinert, T. (1996). A meiotic recombination checkpoint controlled by mitotic checkpoint genes. Nature 383, 840–843.

Modliszewski, J.L., Wang, H., Albright, A.R., Lewis, S.M., Bennett, A.R., Huang, J., Ma, H., Wang, Y., and Copenhaver, G.P. (2018). Elevated temperature increases meiotic crossover frequency via the interfering (Type I) pathway in Arabidopsis thaliana. PLoS Genet 14, e1007384.

Morgan, C.H., Zhang, H., and Bomblies, K. (2017). Are the effects of elevated temperature on meiotic recombination and thermotolerance linked via the axis and synaptonemal complex Philos Trans R Soc Lond B Biol Sci 372.

Muyt, A., Mercier, R., Mezard, C., and Grelon, M. (2009). Meiotic recombination and crossovers in plants. Genome Dyn 5, 14–25.

Nebel, B.R., and Hackett, E.M. (1961). Synaptinemal complexes (cores) in primary spermatocytes of mouse under elevated temperature. Nature 190, 467–468.

Osman, K., Sanchez-Moran, E., Higgins, J.D., Jones, G.H., and Franklin, F.C. (2006). Chromosome synapsis in Arabidopsis: analysis of the transverse filament protein ZYP1 reveals novel functions for the synaptonemal complex. Chromosoma 115, 212–219.

Pacheco, S., Marcet-Ortega, M., Lange, J., Jasin, M., Keeney, S., and Roig, I. (2015). The ATM signaling cascade promotes recombination-dependent pachytene arrest in mouse spermatocytes. PLoS Genet 11, e1005017.

Pao, W.K., and Li, H.W. (1948). Desynapsis and other abnormalities induced by high temperature. J Genet 48, 297–310.

Pecrix, Y., Rallo, G., Folzer, H., Cigna, M., Gudin, S., and Le Bris, M. (2011). Polyploidization mechanisms: temperature environment can induce diploid gamete formation in Rosa sp. J Exp Bot 62, 3587–3597.

Penedos, A., Johnson, A.L., Strong, E., Goldman, A.S., Carballo, J.A., and Cha, R.S. (2015). Essential and Checkpoint Functions of Budding Yeast ATM and ATR during Meiotic Prophase Are Facilitated by Differential Phosphorylation of a Meiotic Adaptor Protein, Hop1. PLoS One 10, e0134297.

Phillips, D., Jenkins, G., Macaulay, M., Nibau, C., Wnetrzak, J., Fallding, D., Colas, I., Oakey, H., Waugh, R., and Ramsay, L. (2015). The effect of temperature on the male and female recombination landscape of barley. New Phytol 208, 421–429.

Prusicki, M.A., Hamamura, Y., and Schnittger, A. (2020). APractical Guide to Live-Cell Imaging of Meiosis in Arabidopsis. Methods Mol Biol 2061, 3–12.

Prusicki, M.A., Keizer, E.M., van Rosmalen, R.P., Komaki, S., Seifert, F., Muller, K., Wijnker, E., Fleck, C., and Schnittger, A. (2019). Live cell imaging of meiosis in Arabidopsis thaliana. Elife 8.

Pusch, S., Dissmeyer, N., and Schnittger, A. (2011). Bimolecular-fluorescence complementation assay to monitor kinase-substrate interactions in vivo. Methods Mol Biol 779, 245–257.

Rockmill, B., Sym, M., Scherthan, H., and Roeder, G.S. (1995). Roles for two RecAhomologs in promoting meiotic chromosome synapsis. Genes Dev 9, 2684–2695.

Roeder, G.S., and Bailis, J.M. (2000). The pachytene checkpoint. Trends Genet 16, 395–403.

Sanchez-Moran, E., Santos, J.L., Jones, G.H., and Franklin, F.C. (2007). ASY1 mediates AtDMC1-dependent interhomolog recombination during meiosis in Arabidopsis. Genes Dev 21, 2220–2233.

Sofroni, K., Takatsuka, H., Yang, C., Dissmeyer, N., Komaki, S., Hamamura, Y., Bottger, L., Umeda, M., and Schnittger, A. (2020). CDKD-dependent activation of CDKA;1 controls microtubule dynamics and cytokinesis during meiosis. J Cell Biol 219.

Song, P., Jia, Q., Chen, L., Jin, X., Xiao, X., Li, L., Chen, H., Qu, Y., Su, Y., Zhang, W., et al. (2020). Involvement of Arabidopsis phospholipase D delta in regulation of ROS-mediated microtubule organization and stomatal movement upon heat shock. J Exp Bot 71, 6555–6570.

Stacey, N.J., Kuromori, T., Azumi, Y., Roberts, G., Breuer, C., Wada, T., Maxwell, A., Roberts, K., and Sugimoto-Shirasu, K. (2006). Arabidopsis SPO11-2 functions with SPO11-1 in meiotic recombination. Plant J 48, 206–216.

Stefani, A., and Colonna, N. (1996). The Influence of Temperature on Meiosis and Microspores Development in Dasypyrum villosum(L.) P. Candargy. CYTOLOGIA 61, 277–283.

Stronghill, P.E., Azimi, W., and Hasenkampf, C.A. (2014). Anovel method to follow meiotic progression in Arabidopsis using confocal microscopy and 5-ethynyl-2’-deoxyuridine labeling. Plant Methods 10, 33.

Stuart, D., and Wittenberg, C. (1998). CLB5 and CLB6 are required for premeiotic DNAreplication and activation of the meiotic S/M checkpoint. Genes Dev 12, 2698–2710.

Su, S.S., and Modrich, P. (1986). Escherichia coli mutS-encoded protein binds to mismatched DNAbase pairs. Proc Natl Acad Sci U S A 83, 5057–5061.

Tang, S., Wu, M.K.Y., Zhang, R., and Hunter, N. (2015). Pervasive and essential roles of the Top3-Rmi1 decatenase orchestrate recombination and facilitate chromosome segregation in meiosis. Mol Cell 57, 607–621.

Wang, J., Li, D., Shang, F., and Kang, X. (2017). High temperature-induced production of unreduced pollen and its cytological effects in Populus. Sci Rep 7, 5281.

Wijnker, E., Harashima, H., Muller, K., Parra-Nunez, P., de Snoo, C.B., van de Belt, J., Dissmeyer, N., Bayer, M., Pradillo, M., and Schnittger, A. (2019). The Cdk1/Cdk2 homolog CDKA;1 controls the recombination landscape in Arabidopsis. Proc Natl Acad Sci U S A 116, 12534–12539.

Wijnker, E., and Schnittger, A. (2013). Control of the meiotic cell division programin plants. Plant Reprod 26, 143–158.

Wilson, J.Y. (1959). Duration of meiosis in relation to temperature. Heredity 13, 263–267.

Wu, S., Scheible, W.R., Schindelasch, D., Van Den Daele, H., De Veylder, L., and Baskin, T.I. (2010). Aconditional mutation in Arabidopsis thaliana separase induces chromosome non-disjunction, aberrant morphogenesis and cyclin B1;1 stability. Development 137, 953–961.

Yahya, G., Perez, A.P., Mendoza, M.B., Parisi, E., Moreno, D.F., Artes, M.H., Gallego, C., and Aldea, M. (2021). Stress granules display bistable dynamics modulated by Cdk. J Cell Biol 220.

Yang, C., Hamamura, Y., Sofroni, K., Bower, F., Stolze, S.C., Nakagami, H., and Schnittger, A. (2019). SWITCH 1/DYAD is a WINGS APART-LIKE antagonist that maintains sister chromatid cohesion in meiosis. Nat Commun 10, 1755.

Yang, C., Sofroni, K., Wijnker, E., Hamamura, Y., Carstens, L., Harashima, H., Stolze, S.C., Vezon, D., Chelysheva, L., Orban-Nemeth, Z., et al. (2020). The Arabidopsis Cdk1/Cdk2 homolog CDKA;1 controls chromosome axis assembly during plant meiosis. EMBO J 39, e101625.

Yao, Y., Li, X., Chen, W., Liu, H., Mi, L., Ren, D., Mo, A., and Lu, P. (2020). ATM Promotes RAD51-Mediated Meiotic DSB Repair by Inter- Sister-Chromatid Recombination in Arabidopsis. Front Plant Sci 11, 839.

Yazawa, T., Nakayama, Y., Fujimoto, K., Matsuda, Y., Abe, K., Kitano, T., Abe, S., and Yamamoto, T. (2003). Abnormal spermatogenesis at low temperatures in the Japanese red-bellied newt, Cynops pyrrhogaster: possible biological significance of the cessation of spermatocytogenesis. Mol Reprod Dev 66, 60–66.

Yue, Y., Zhang, P., and Shang, Y. (2019). The potential global distribution and dynamics of wheat under multiple climate change scenarios. Sci Total Environ 688, 1308–1318.

Zhao, X., Bramsiepe, J., Van Durme, M., Komaki, S., Prusicki, M.A., Maruyama, D., Forner, J., Medzihradszky, A., Wijnker, E., Harashima, H., et al. (2017). RETINOBLASTOMA RELATED1 mediates germline entry in Arabidopsis. Science 356.

Zhao, X., Harashima, H., Dissmeyer, N., Pusch, S., Weimer, A.K., Bramsiepe, J., Bouyer, D., Rademacher, S., Nowack, M.K., Novak, B., et al. (2012). Ageneral G1/S-phase cell-cycle control module in the flowering plant Arabidopsis thaliana. PLoS Genet 8, e1002847.

Zuela, N., and Gruenbaum, Y. (2016). Matefin/SUN-1 Phosphorylation on Serine 43 Is Mediated by CDK-1 and Required for Its Localization to Centrosomes and Normal Mitosis in C. elegans Embryos. Cells 5.

